# Robust and fast Monte Carlo Markov Chain sampling of diffusion MRI microstructure models

**DOI:** 10.1101/328427

**Authors:** R.L. Harms, A. Roebroeck

**Affiliations:** Dept. of Cognitive Neuroscience, Faculty of Psychology & Neuroscience, Maastricht University, the Netherlands

**Keywords:** Monte Carlo Markov Chain (MCMC) sampling, Diffusion MRI, Microstructure, Biophysical compartment models, Parallel computing, GPU computing

## Abstract

In diffusion MRI analysis, advances in biophysical multi-compartment modeling have gained popularity over the conventional Diffusion Tensor Imaging (DTI), because they possess greater specificity in relating the dMRI signal to underlying cellular microstructure. Biophysical multi-compartment models require parameter estimation, typically performed using either Maximum Likelihood Estimation (MLE) or using Monte Carlo Markov Chain (MCMC) sampling. Whereas MLE provides only a point estimate of the fitted model parameters, MCMC recovers the entire posterior distribution of the model parameters given the data, providing additional information such as parameter uncertainty and correlations. MCMC sampling is currently not routinely applied in dMRI microstructure modeling because it requires adjustments and tuning specific to each model, particularly in the choice of proposal distributions, burn-in length, thinning and the number of samples to store. In addition, sampling often takes at least an order of magnitude more time than non-linear optimization. Here we investigate the performance of MCMC algorithm variations over multiple popular diffusion microstructure models to see whether a single well performing variation could be applied efficiently and robustly to many models. Using an efficient GPU-based implementation, we show that run times can be removed as a prohibitive constraint for sampling of diffusion multi-compartment models. Using this implementation, we investigated the effectiveness of different adaptive MCMC algorithms, burn-in, initialization and thinning. Finally we apply the theory of Effective Sample Size to diffusion multi-compartment models as a way of determining a relatively general target for the number of samples needed to characterize parameter distributions for different models and datasets. We conclude that robust and fast sampling is achieved in most diffusion microstructure models with the Adaptive Metropolis-Within-Gibbs (AMWG) algorithm initialized with an MLE point estimate, in which case 100 to 200 samples are sufficient as a burn-in and thinning is mostly unnecessary. As a relatively general target for the number of samples, we recommend a multivariate Effective Sample Size of 2200.

## 1 Introduction

Advances in microstructure modeling of diffusion Magnetic Resonance Imaging (dMRI) data have recently gained popularity since they possess greater specificity than Diffusion Tensor Imaging (DTI) in relating the dMRI signal to the underlying cellular microstructure, such as axonal density, orientation dispersion or diameter distributions. Typically, dMRI models are fitted to the data using non-linear optimization (Assaf et al., 2004; Assaf & Basser, 2005; Assaf et al., 2008; Panagiotaki et al., 2012; Zhang et al., 2012; Assaf et al., 2013; Fieremans et al., 2013; De Santis et al., 2014b,a; Jelescu et al., 2015b; Harms et al., 2017) or linear convex optimization (Daducci et al., 2015) methods to obtain a parameter point estimate per voxel. These point estimates provide scalar maps over the brain of micro-structural parameters, such as the fraction of restricted diffusion as a proxy for fiber density. These point estimates however do not include the uncertainty in the estimate, nor do they include the interdependency of parameters. The gold standard of obtaining these quantities is by using Monte Carlo Markov Chain (MCMC) sampling, as for example in (Behrens et al., 2003; Alexander, 2008; Alexander et al., 2010; Sotiropoulos et al., 2013). MCMC generates, per voxel, a multi-dimensional chain of samples, the stationary distribution of which is the posterior distribution, i.e. the probability density of the model parameters given the data. Per voxel, these samples capture parameter dependencies, multimodality and the width of peaks around optimal parameter values. For instance, summarizing the chain under Gaussian assumptions with a sample covariance matrix, would already provide mean parameter estimates and corresponding uncertainties (the standard deviation), as well as inter-parameter correlations (Figure 1).

Despite these advantages, MCMC sampling is currently not routinely applied in dMRI microstructure modeling since it often requires adjustments and tuning specific to each model, particularly in the choice of proposals, burn-in length, thinning and the number of samples to store. In addition, sampling often takes at least an order of magnitude more time than nonlinear optimization.

Here we investigate the performance of a few variants of Random Walk Metropolis MCMC algorithms over multiple popular diffusion microstructure models to see whether a single well performing variation could be applied efficiently and robustly to many models. To this end, we evaluate different strategies for adaptive proposals, burn-in and thinning with respect to diffusion MRI modeling. To determine a lower bound on the number of samples needed, we apply the concept of effective sample sizes to determine information content and posterior confidence. Finally, to reduce run-time constraints we provide an efficient parallel GPU implementation of all models and MCMC algorithms in the open source Microstructure Diffusion Toolbox (MDT; https://github.com/cbclab/MDT).

**Figure 1:**
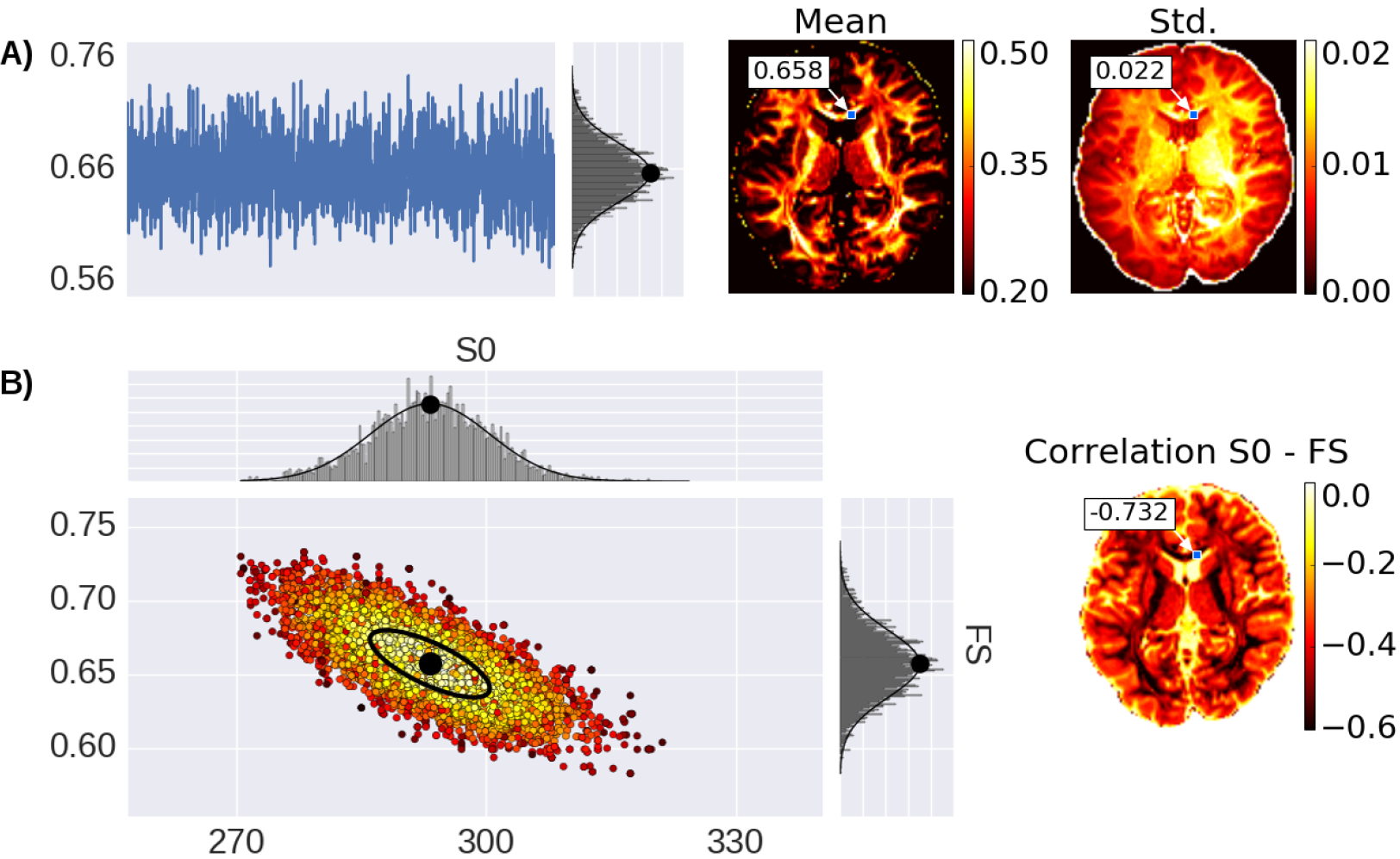
Illustration of parameter uncertainty and correlation for the Ball&Stick model using MCMC sampling, with the Fraction of Stick (FS) and the non-diffusion weighted signal intensity (S0). **A)** On the left, a single FS sampling trace and its corresponding histogram for the highlighted voxel with a Gaussian distribution function fitted to the samples with its mean indicated by a black dot. On the right, the mean and standard deviation (std.) maps generated from the independent voxel chains per voxel. **B)** On the left, the scatter-plot for two parameters (FS and S0) with the corresponding marginal histograms for the voxel highlighted in the maps. On the right, the S0-FS correlation map.

## 2 Methods

The biophysical (multi-)compartment models and the Markov Chain Monte Carlo (MCMC) algorithms used in this study are implemented in a Python based GPU accelerated toolbox (the Microstructure Diffusion Toolbox, MDT, freely available under an open source L-GPL license at https://github.com/cbclab/MDT). Its modular design allows arbitrary combinations of models with likelihood and prior distributions. The MCMC implementations are voxel-wise parallelized using the OpenCL framework, allowing parallel computations on multi-core CPU and/or Graphics Processing Units (GPUs).

We use the models and MCMC routine as implemented in MDT version 0.10.9. Unless stated otherwise, we initialize the MCMC sampling with a Maximum Likelihood Estimator (MLE) obtained from non-linear parameter optimization using the Powell routine with cascaded model initialization (Harms et al., 2017).

First, we define and review posteriors, likelihoods and priors relevant to diffusion multi-compartment models. We next define the Metropolis-Hastings as the general type of Markov Chain Monte Carlo algorithms used in this work. Then, under assumptions of symmetric and current position centered proposals, updated one dimension at a time, we derive the Metropolis-Within-Gibbs algorithm. The Metropolis-Within-Gibbs algorithm is then explained with and without the use of adaptive proposals. We subsequently define burn-in, thinning, effective sample size and number of samples as the targets of investigation for diffusion microstructure models.

### 2.1 Posterior, likelihoods and priors

Given observations 𝒪 and a model with parameters **x** ∈ ℝ^*n*^, we can construct a posterior distribution p(**x**|𝒪) from a log-likelihood distribution *l*(𝒪|**x**) and prior distribution *p*(**x**), as:

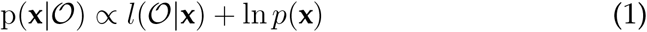

In this work we are interested in approximating the posterior density of: p(**x**|𝒪) using MCMC sampling.

#### 2.1.1 Likelihood distribution

The likelihood distribution *l*(𝒪|**x**) contains a signal model, embedding the diffusion microstructure modeling assumptions combined with a noise model. As discussed in previous work (Harms et al., 2017; Panagiotaki et al., 2012; Alexander, 2009), we use the Offset Gaussian model as likelihood distribution:

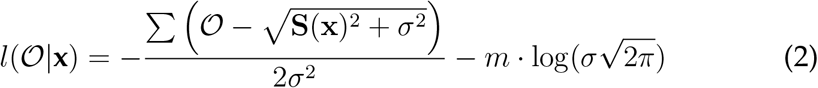

with *l*(𝒪|**x**) the log-likelihood function, **x** the parameter vector, O the observations (the data volumes), **S**(**x**) the signal model, *σ* the standard deviation of the Gaussian distributed noise (of the complex valued data, i.e. before calculation of magnitude data) and *m* the number of volumes in the dataset (number of observations). We estimated *σ* a priori from the reconstructed magnitude images using the *σ*_mult_ method in (Dietrich et al., 2007, eq. A6). For signal model naming we use the postfix ‘_in[n]’ to identify the number of restricted compartments employed in models which allow multiple intra-axonal compartments (Harms et al., 2017). For example, CHARMED_in2 indicates a CHARMED model with 2 intra-axonal compartments (and the regular single extra-axonal compartment), for each of two unique fiber orientations in a voxel.

#### 2.1.2 Priors

The prior distribution *p*(**x**) describes the a priori knowledge we have about the model and its parameters. We construct a complete model prior as a product of priors per parameter, *p*_*i*_(**x**_*i*_) (see table 1), with one or more model specific priors over multiple parameters, *p*_*j*_(**x**|*M*), for model prior *j* of model *M* (see table 2):

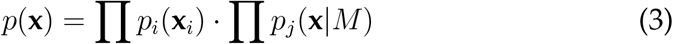

Assuming no further a priori knowledge than logical or biologically plausible ranges, we use uniform priors for each parameter, *p*_*i*_(**x**_*j*_) ~ *U*(*a*,*b*). Additionally, for multi-compartment models with volume fraction weighted compartments (i.e. Ball&Stick_in1, NODDI and CHARMED_in1) we add a prior on the *n* − 1 volume fractions *w*_*k*_ to ensure 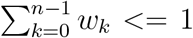, to ensure proper volume fraction weighting. Note that the last volume fraction is not sampled but is set to one minus the sum of the others, 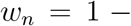 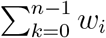. To the Tensor compartment (used in the Tensor and CHARMED_-in1 model), we add a prior to ensure strictly decreasing diffusivities 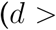 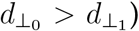, this prevents parameter aliasing of the Tensor orientation parameters (see (Gelman et al., 2013) on aliasing).

**Table 1:** The priors *p*_*i*_(**x**_*i*_) per model parameter. These priors are combined with the model specific parameters in table 2 to form the complete model priors. For parameter usage and specification see (Harms et al., 2017).

| Parameter | Prior | Used in model(s) |
| --- | --- | --- |
| $S_0$ | $U(0, 1 \cdot 10^{10})$ | Tensor, Ball&Stick_in1, NODDI, CHARMED_in1 |
| $w_i$ | $U(0, 1)$ | Ball&Stick_in1, NODDI, CHARMED_in1 |
| $d_{\parallel}$ (or $d$) | $U(3 \cdot 10^{-11}, 1 \cdot 10^{-8})$ | Tensor, Ball&Stick_in1, NODDI, CHARMED_in1 |
| $d_{\perp_1}, d_{\perp_2}$ | $U(0, 1 \cdot 10^{-8})$ | Tensor, CHARMED_in1 |
| $\theta, \phi$ | $U(0, \pi)$ | Tensor, Ball&Stick_in1, NODDI, CHARMED_in1 |
| $\psi$ | $U(0, \pi)$ | Tensor |
| $\kappa$ | $U(0, 2\pi)$ | NODDI |

**Table 2:**
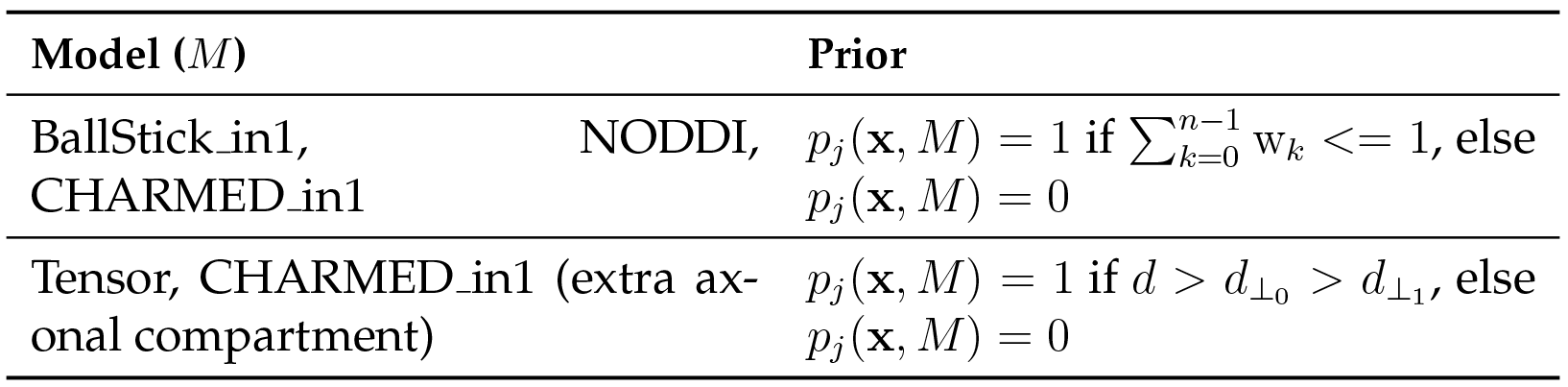
The model priors *p*_*j*_ (**x**|*M*) for model *M*. Each of these priors should be interpreted as a boolean, that is, they return a value of 1 if the condition is fulfilled, else they return 0. These priors are combined with the parameter specific priors in table 1 to form the complete model priors.

| Model ($M$) | Prior |
| --- | --- |
| BallStick_in1, NODDI, CHARMED_in1 | $p_j(\mathbf{x}, M) = 1$ if $\sum_{k=0}^{n-1} w_k \leq 1$, else $p_j(\mathbf{x}, M) = 0$ |
| Tensor, CHARMED_in1 (extra axonal compartment) | $p_j(\mathbf{x}, M) = 1$ if $d > d_{\perp_0} > d_{\perp_1}$, else $p_j(\mathbf{x}, M) = 0$ |

### 2.2 Markov Chain Monte Carlo

Markov Chain Monte Carlo (MCMC) is a class of numerical approximation algorithms for sampling from the probability density function *π*(·) of a target random variate, by generating a Markov chain **X**^(0)^, **X**^(1)^,… with stationary distribution *π*(·). There are a large number of MCMC algorithms, including Metropolis-Within-Gibbs (a.k.a Metropolis) (Metropolis et al., 1953), Metropolis-Hastings (Hastings, 1970), Gibbs (Turchin, 1971; Geman & Geman, 1984), Component-wise Hit-And-Run Metropolis (Turchin, 1971; Smith, 1984), Random Walk Metropolis (Muller, 1994), Multiple-Try Metropolis (Liu et al., 2000), No-U-Turn sampler (Hoffman & Gelman, 2011) and many more. All of these algorithms are known as special cases of the Metropolis-Hastings algorithm and differ only in the proposal distributions they employ (Johnson et al., 2013; Chib & Greenberg, 1995).

The general Metropolis-Hastings algorithm works as follows. Given a current position **X**^(*t*)^ at step *t* on a *p*-dimensional Markov chain, a new position **X**^(*t*+1)^ is obtained by generating a candidate position **Y** from the proposal density *q*(**X**^(*t*)^|·), which is then either accepted with probability *α*, or rejected with probability 1 − *α*. If the candidate position is accepted, **X**^(*t*+1)^ = **y**, else, **X**^(*t*+1)^ = **X**^(*t*)^. The acceptance criteria *α* is a function given by (Hastings, 1970):

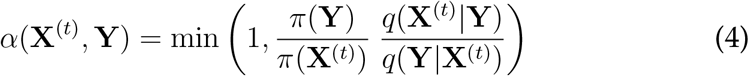

where *π*(·) is our target density, generally given by our posterior distribution function p(**X**|·). The subsequent collection of points {**X**^(0)^,…, **X**^(*s*)^} for a sample size s is called the chain and is the algorithm’s output. The *ergodic* property of this algorithm guarantees that this chain converges (in the long run) to a stationary distribution which approximates the target density function *π*(·) (Metropolis et al., 1953; Hastings, 1970).

In this work we use a symmetric proposal distribution centered around the current sampling position for every dimension (every component) of the sampled multivariate distribution. If the proposal distribution *q* is symmetric, i.e. *q*(**X**^(*t*)^|**Y**) = *q*(**Y**|**X**^(*t*)^) we can drop Hasting’s addition to the acceptance criteria function, simplifying it to the Metropolis criteria (Metropolis et al., 1953):

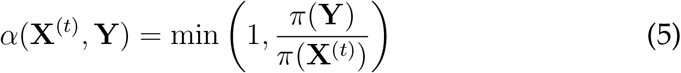

If furthermore *q*(**X**^(*t*)^|**Y**) = *q*(**X**^(*t*)^ − **Y**) = *q*(**Y** − **X**^(*t*)^), that is, the proposal is centered around **X**^(*t*)^ ∀*t*, we typically denote it as the Random Walk Metropolis algorithm (Robert, 2015; Chib & Greenberg, 1995; Johnson et al., 2013; Sherlock et al., 2010; Muller, 1994; Hastings, 1970; Metropolis et al., 1953)..

#### 2.2.1 Metropolis-Within-Gibbs

In the component wise updating scheme, a new position **X**^(*t*+1)^ is proposed one component (i.e. one dimension) at a time, in contrast to updating all *p* dimensions at once. Since such a single component updating Random Walk Metropolis algorithm uses elements both of Gibbs sampling and of Metropolis-Hastings, this scheme is also referred to as Metropolis-Within-Gibbs (MWG) (van Ravenzwaaij et al., 2016; Robert, 2015; Sherlock et al., 2010). Let 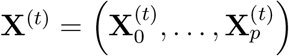 define the components 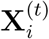 of **X**^(*t*)^, then we can define

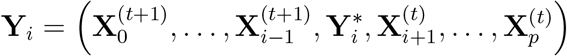

as the candidate position for component *i*, and

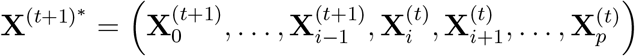

as the temporary position in the chain while component *i* is being updated. The proposals 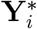 are generated using the symmetric proposal *q*_*i*_(**X**^(*t*+1)^*|·) which updates the *i*th component dependent on the components already updated. One iteration of the MWG algorithm cycles through all *i* components, where each proposal **Y**_*i*_ is accepted or rejected using probability *α*(**X**^(*t*+1)^*, **Y**_*i*_).

### 2.3 Proposal distributions

As symmetric proposal distributions for our MWG algorithm we use centered Normal distributions, i.e. 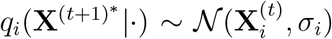, where *σ*_*i*_ is the proposal standard deviation of the *i*th component (not to be confused with the *σ* used in the likelihood distribution above). For the orientation parameters *θ*, *ϕ* and *ψ* we use a circular Normal modulus *π*, i.e. 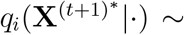 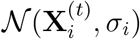 mod *π*. See table 3 for an overview of the default proposal distributions used per parameter.

**Table 3:** The proposal distributions *q*_*i*_(**X**^(*t*+1)^*|·) per model parameter with their default proposal standard deviations. For parameter usage and specification see (Harms et al., 2017).

| Parameter | Prior |
| --- | --- |
| $S_0$ | $\mathcal{N}(\mathbf{X}_i^{(t)}, 10)$ |
| $w_i$ | $\mathcal{N}(\mathbf{X}_i^{(t)}, 0.01)$ |
| $d_{\parallel}$ (or $d$), $d_{\perp_1}$, $d_{\perp_2}$ | $\mathcal{N}(\mathbf{X}_i^{(t)}, 1 \cdot 10^{-8})$ |
| $\theta, \phi, \psi$ | $\mathcal{N}(\mathbf{X}_i^{(t)}, 0.1) \bmod \pi$ |
| $\kappa$ | $\mathcal{N}(\mathbf{X}_i^{(t)}, 0.01)$ |

### 2.4 Adaptive Metropolis

While in the traditional Metropolis-Within-Gibbs algorithm each *σ*_*i*_ in the proposal distribution is fixed, variations of this algorithm exist that autotune each *σ*_i_ to improve the information content of the Markov chain. While technically each of these variations is a distinct MCMC algorithm, we consider and compare three of these variations here as proposal updating strategies for the MWG algorithm.

The first adaptation strategy compared is the Single Component Adaptive Metropolis (*SCAM*) algorithm (Haario et al., 2005), which works by adapting the proposal standard deviation to the empirical standard deviation of the component’s marginal distribution. That is, the standard deviation 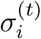 for the proposal distribution of the *i*th component at time *t* is given by:

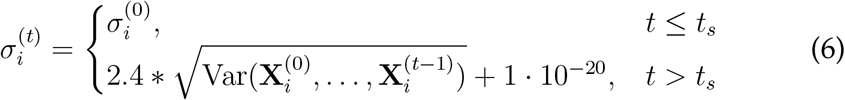

where *t*_*s*_ denotes the iteration after which the adaptation starts (we use *t*_*s*_ = 100). A small constant is necessary to prevent the standard deviation from shrinking to zero. This adaptation algorithm has been proven to retain ergodicity, meaning it is guaranteed to converge to the right stationary distribution (Haario et al., 2005).

The other two methods work by adapting the acceptance rate of the generated proposals. The acceptance rate is the ratio of accepted to generated proposals and is typically updated batch-wise. In general, by decreasing the proposal standard deviation the acceptance rate increases and vice versa. Theoretically, for single component updating schemes (like in this work), the optimal target acceptance rate is 0.44 (Gelman et al., 1996).

The first of the two acceptance rate scaling strategies is from the FSL Bed-postX software package. This strategy, which we refer to as the *FSL* strategy, tunes the acceptance rate to 0.5 (Behrens et al., 2003). It works by multiplying the proposal variance by the ratio of accepted to rejected samples, i.e. it multiplies the standard deviation *σ*_*i*_ by 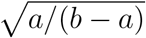 after every batch of size *b* with *a* accepted samples. We update the proposals after every batch of size 50 (*b* = 50) (Behrens et al., 2003). Since this method never ceases the adaptation of the standard deviations, it theoretically loses er-godicity of the chain (Roberts & Rosenthal, 2009, 2007).

The last method, the Adaptive Metropolis-Within-Gibbs (*AMWG*) method (Roberts & Rosenthal, 2009) uses the current acceptance rate over batches to tune the acceptance rate to 0.44. After the *n*th batch of 50 iterations (Roberts & Rosenthal, 2009), this method updates the logarithm of o_i_ by adding or subtracting an adoption amount 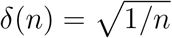 depending on the acceptance rate of that batch. That is, after every batch, *σ*_*i*_ is updated by:

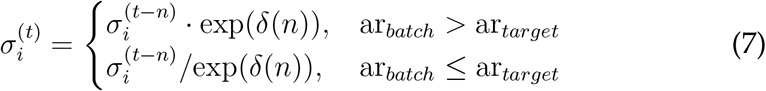

where ar_*batch*_ is the acceptance rate of the current batch and ar_*target*_ is the target acceptance rate (0.44). Since this method features diminishing adaptation, the chain remains ergodic (Roberts & Rosenthal, 2009).

We compare all three strategies and the default, no adaptation, on the number of effective samples they generate (see below) and on accuracy and precision using ground truth simulation data. We sample all models with 10000 samples, without thinning, using the point optimized Maximum Likelihood Estimator (MLE) as starting point and with a burn-in of 1000. Estimates of the standard error of the mean (SEM) are obtained by averaging the statistics over 10 independent MCMC runs.

### 2.5 Burn-in

Burn-in is the process of discarding the first *z* samples from the chain and using only the remaining samples in subsequent analysis. The idea is that if the starting point had a low probability then the limited number of early samples may oversample low probability regions. By discarding the first *z* samples as a burn-in, the hope is that, by then, the chain has converged to its stationary distribution and that all further samples are directly from the stationary distribution (Robert, 2015). Theoretically, burn-in is unnecessary since any empirical average

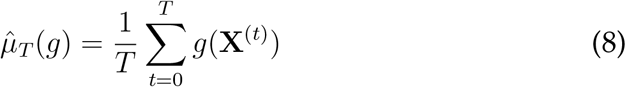

for any function *g* will convert to *μ*(*g*) given a large enough sample size and given that the chain is ergodic (Robert, 2015). Additionally, since it can not be predicted how long it will take for the chain to reach convergence, the required burn-in can only be estimated post-hoc. In practice, discarding the first few thousand samples as a burn-in often works and is less time-consuming than generating a lot of samples to average out the effects of a low probability starting position.

An alternative to burn-in, or, to reduce the need for burn-in, is to use a Maximum Likelihood Estimator as starting point for the MCMC sampling (van Ravenzwaaij et al., 2016). If the optimization routine did its work well, the MLE should be part of the stationary distribution of the Markov chain, removing the need for burn-in altogether. We compare initialization using a MLE obtained using the Powell routine (Harms et al., 2017), with a initialization from a default a priori value (table 4). For most models the MLE optimization results can be used directly, for the Tensor model we sometimes need to sort the diffusivities and reorient the *θ*, *ϕ* and *ψ* angles to ensure decreasing diffusivities. To evaluate the effect of burn-in and initialization single-slice datas was sampled using the NODDI model with the default starting point and with MLE. For selected single voxels the NODDI model was also sampled using the MLE starting point and two random volume fractions as a starting point.

### 2.6 Thinning

Thinning is the process of using only every *k*th step of the chain for analysis, while all other steps are discarded, with as goal reducing autocorrelation and obtaining relatively independent samples. Several authors have recommended against the use of thinning, stating that it is often unnecessary, always inefficient and reduces the precision of the posterior estimates (Link & Eaton, 2012; MacEachern & Berliner, 1994; Jackman, 2009; Geyer, 1991; Christensen et al., 2010).

**Table 4:** The default starting points for the MCMC sampler, used only when the sampler is not initialized with a maximum likelihood estimator.

| Parameter | Default starting value |
| --- | --- |
| $S_0$ | $1 \cdot 10^4$ |
| $w_i$ | 0.5 |
| $d_{\parallel}$ (or $d$) | $1.7 \cdot 10^{-9}$ |
| $d_{\perp_1}$ | $1.7 \cdot 10^{-10}$ |
| $d_{\perp_2}$ | $1.7 \cdot 10^{-11}$ |
| $\theta, \phi, \psi$ | $\pi/2$ |
| $\kappa$ | 1 |

The only valid reason for thinning is to avoid bias in the standard error estimate of posterior mean, when that mean estimate was computed over all (non thinned) samples (Link & Eaton, 2012; MacEachern &; Berliner, 1994). In general, thinning is only considered worthwhile if there are storage limitations, or when the cost of processing the output outweighs the benefits of reduced variance of the estimator (Geyer, 1991; MacEachern & Berliner, 1994; Link & Eaton, 2012).

To evaluate the effect of thinning we sampled a single voxel with 20000 samples and compared the effect of using all samples in computing the posterior mean and posterior standard deviation of a volume fraction against using only a thinned amount of samples We compare the effect of taking *n* samples with a thinning of *k* (the *thinning* method) against just using all *n* · *k* samples (the *more samples* method).

### 2.7 Effective Sample Size

The Effective Sample Size (ESS) in the context of MCMC, measures the information content, or effectiveness of a sample chain. For example, 1000 samples with an ESS of 200 have a higher information content than 2000 samples with an ESS of 100. The ESS can be defined as the minimum size of a set of posterior samples (taken directly from the posterior), which have the same efficiency (measure of quality) in the posterior density estimation as a given chain of samples obtained from MCMC sampling (Martino et al., 2017). Conversely, ESS theory can quantify how many samples should be taken in a chain to reach a given quality of posterior estimates. We use the ESS theory to comparing proposal adaptation strategies and to estimating the minimum number of samples necessary for adequate sampling of diffusion microstructure models.

Multivariate ESS theory (Vats et al., 2015) is an extension of univariate ESS theory (Gong & Flegal, 2016; Liu, 2004; Robert & Casella, 2004; Kass et al., 1998) and computes the empirical ESS as:

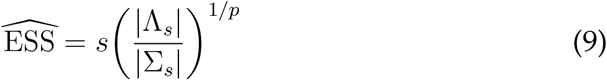

with *s* is the number of obtained samples, *p* the number of parameters, Λ_*s*_ the covariance matrix of the samples and ∑_*s*_ an estimate of the Monte Carlo standard error (the error in the chain caused by the MCMC sampling process), here calculated using a batch means algorithm (Vats et al., 2015).

### 2.8 Number of samples

The multivariate ESS theory dictates that one can terminate the sampling when the empirical number of effective samples, 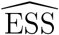, satisfies:

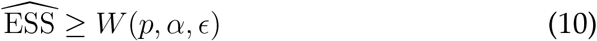

where *W*(*p*, *α*, *ϵ*) gives a theoretical lower bound with *p* the number of parameters in the model, *α* the level of confidence of a desired confidence region and *ϵ* a desired relative precision (the relative contribution of Monte Carlo error to the variability in the target distribution). *W*(*p*, *α*, *ϵ*) can be determined a priori and is defined as:

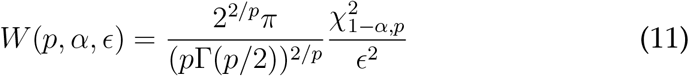

with *χ*^2^ the chi-square function and Γ(·) the Gamma function (Vats et al., 2015). Figure 2 shows the effect of *α* and *ϵ* on *W*(*p*, *α*, *ϵ*). Given the exponential increase in the number of samples need for very high confidence and precision, we aim for a 95% confidence region (*α* = 0.05) with a 90% precision (*ϵ* = 0.1) in this work.

Since online monitoring of the ESS (during MCMC sampling) is an expensive operation, and terminating on ESS will yield different sample sizes for different voxels, we instead use the ESS theory to estimate a fixed minimum number of samples needed to reach a desired ESS when averaged over a white matter mask. We sampled with the BallStick_in1, Tensor, NODDI and CHARMED_in1 models, using respectively 15000, 20000, 20000 and 30000 samples and computed from those samples the average ESS over white matter masks. For *α* = 0.05 and *ϵ* = 0.1 we computed per model the theoretical minimum required effective sample size *W*(*p*, *α*, *ϵ*). We compared those theoretical numbers to the obtained average effective sample size and estimated a minimum required number of samples *ŝ* using the ratio:

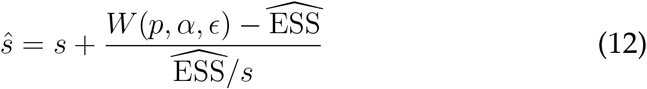

where *s* is the number of samples we started out with, *W*(*p*, *α*, *ϵ*) the theoretical ESS requirements and 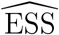 the estimated number of effective samples in our chain when averaged over the white matter mask. As an estimate of computation times, we record runtime statistics for sampling the recommended number of samples using an AMD Fury X graphics card.

### 2.9 Datasets

For this study we used two groups of ten subjects coming from two studies, each whith a different acquisition protocol. The first ten subjects are from the freely available fully preprocessed dMRI data from the USC-Harvard consortium of the Human Connectome project. Data used in the preparation of this work were obtained from the MGH-USC Human Connectome Project (HCP) database (https://ida.loni.usc.edu/loginjsp). The data were acquired on a specialized Siemens Magnetom Connectom with 300mT/m gradient set (Siemens, Erlangen, Germany). These datasets were acquired at a resolution of 1.5mm isotropic with Δ=21.8ms, *δ*=12.9ms, TE=57ms, TR=8800ms, Partial Fourier = 6/8, MB factor 1 (i.e. no simultaneous multi-slice), in-plane GRAPPA acceleration factor 3, with 4 shells of b=1000, 3000, 5000, 10,000 s/mm^2^, with respectively 64, 64, 128, 393 directions to which are added 40 interleaved b0 volumes leading to 552 volumes in total per subject, with an acquisition time of 89 minutes. We refer to these datasets as *HCP MGH − 1.5mm −552vol − b10k* and to the multi-shell direction table as the *HCP MGH* table. These four-shell, high number of directions, and very high maximum b- value datasets allow a wide range of models to be fitted.

**Figure 2:**
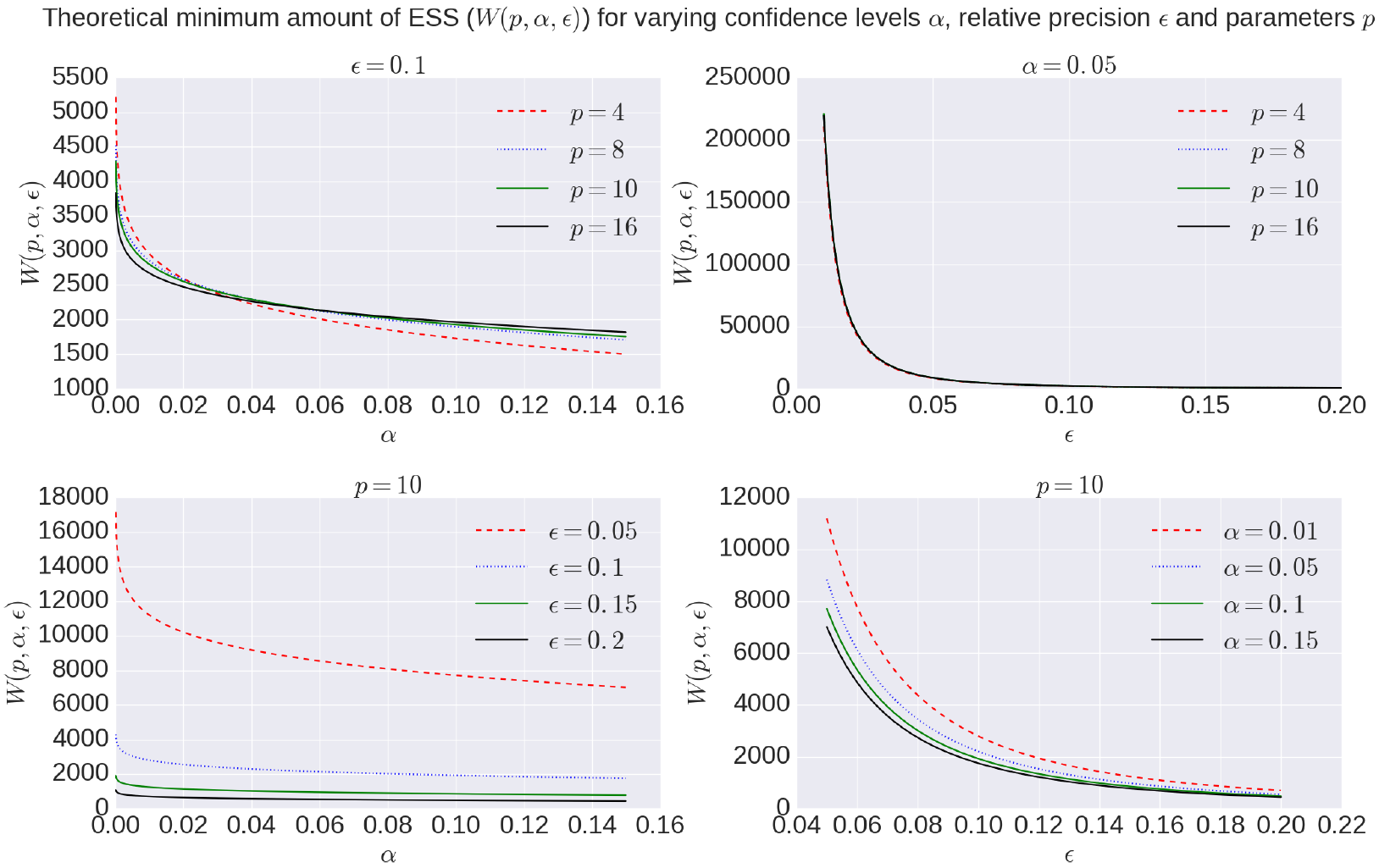
Overview of theoretical minimum ESS, *W*(*p*, *α*, *ϵ*), to reach a specific confidence level *α* with a desired relative precision *ϵ* for a model with number of parameters *p*.

The second set of ten subjects comes from the diffusion protocol pilot phase of the Rhineland Study (www.rheinland-studie.de) and was acquired on a Siemens Magnetom Prisma (Siemens, Erlangen, Germany) with the Center for Magnetic Resonance Research (CMRR) multi-band (MB) diffusion sequence (Moeller et al., 2010; Xu et al., 2013). These datasets had a resolution of 2.0mm isotropic with Δ=45.8ms, *δ*=16.3ms and TE=90ms, TR=4500ms Partial Fourier = 6/8, MB factor 3, no in-plane acceleration with 3 shells of b=1000, 2000, 3000 s/mm^2^, with respectively 30, 40 and 50 directions to which are added 14 interleaved b0 volumes leading to 134 volumes in total per subject, with an acquisition time of 10 min 21 sec. Additional b0 volumes were acquired with a reversed phase encoding direction which were used to correct susceptibility related distortion (in addition to bulk subject motion) with the topup and eddy tools in FSL version 5.0.9. We refer to these datasets as *RLS − 2mm − 134dir − b3k* and to the multi-shell direction table as the RLS table. These three-shell datasets represent a relatively short time acquisition protocol that still allows many models to be fitted.

Since the Tensor model is only valid for b-values up to about 1200s/mm^2^, we only use the b-value 1000s/mm^2^ shell and b0 volumes during model optimization and sampling. All other models are estimated on all data volumes. For all datasets we created a white matter (WM) mask and, using BET from FSL (Smith, 2002), a whole brain mask. The whole brain mask is used during sampling, whereas averages over the WM mask are used in model or data comparisons. The WM mask was calculated by applying a lower threshold of 0.3 on the Tensor FA results, followed by a double pass 3D median filter of radius 2 in all directions. The Tensor estimate for this mask generation was calculated using a CI Ball Stick/Tensor cascade optimized with the Powell method (Harms et al., 2017).

### 2.10 Ground truth simulations

We performed ground truth simulations to illustrate the effects of the adaptive proposals on the accuracy and precision of parameter estimation. For all models in the study, we simulated 10000 repeats with random volume fractions, diffusivities and orientations, using both a HCP MGH and a RLS multi-shell direction table with Rician noise of an SNR of 30. For the Tensor model we only use the b-value 1000s/mm^2^ shell and b0 volumes of the acquisition tables. To ensure Gaussianity of the sampled parameter distributions, we generate the parameters with a smaller range than the support of the sampling priors (table 5). To allow an uniform SNR of 30 we fix *S*_0_ to 1 · 10^4^.

Analogous to (Harms et al., 2017), we compute estimation error as the mean of the (marginal) posterior minus ground truth parameter value for the intra-axonal volume fraction, i.e. fraction of stick (FS) for Ball&Sticks_-in1, fraction of restricted (FR) for CHARMED_in1 and fraction of restricted (FR) for NODDI. We compute a measure of accuracy as the inverse of the mean of the average estimate error over ten thousand random repeats and a measure of precision as the inverse of the standard deviation of the average estimates. Finally, we aggregate these results per model and per experiment over 10 independent ground truth simulation trials into a mean and standard error of the mean (SEM) for both accuracy and precision. When reported, the effective sample size (ESS) is computed using the multivariate ESS theory, averaged over the 10000 voxels with again a SEM over 10 trials.

**Table 5:**
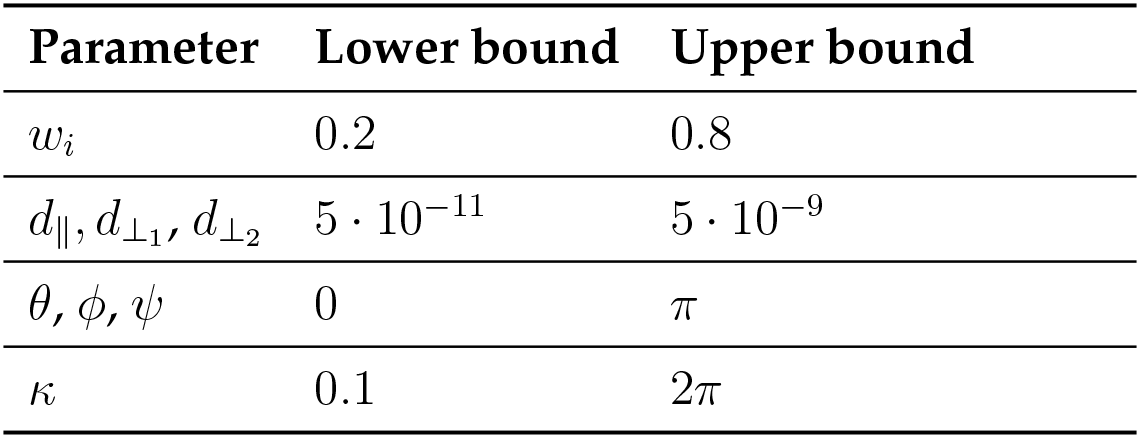
The simulation ranges per model parameters. We generate uniformly distributed parameter values using the upper and lower bounds presented.

## 3 Results

We begin by comparing the four different proposal strategies for sampling the different microstructure compartment models: Tensor, Ball&Sticks_-in1, CHARMED_in1 and NODDI. We then present burn-in and thinning given an effective proposal strategy, and end with ESS estimates on the minimum number of samples needed for adequate characterization of the posterior distribution.

### 3.1 Adaptive proposal strategies

We compare three different adaptive proposal strategies, the Single Component Adaptive Metropolis (SCAM), the FSL acceptance rate scaling (FSL) and the Adaptive Metropolis-Within-Gibbs (AMWG), against the default of no adaptive proposals (None). Comparisons are based on multivariate Effective Sample Size, and accuracy and precision using ground truth simulations. Figure 3 illustrates the effect of using MCMC algorithms with adaptive proposal strategies using the Ball&Stick_in1 model, the HCP MGH dataset, an initial standard deviation of 0.25, after a burn-in of 1000 steps. The illustration clearly shows that without adaptive proposals the chain can get stuck in the same position for quite some time, while all adaptive proposal methods can adapt the standard deviations to better cover the support of the posterior distribution.

The empirical ESS (eq. 9) measures the information content or effectiveness of a sample chain. As such, comparing the ESS for an equal number of actual samples for different proposal strategies evaluates how effectively each strategy generates useful information about the posterior distribution. Figure 4 shows that all adaptive methods clearly outperform the default, None, by generating about 2~3 times more effective samples for equal length chains. The AMWG method generates the largest ESS in all cases, although with a small margin compared to the other two adaptive methods. Compared on accuracy and precision in ground truth simulations (figure 5), the adaptive proposal methods again show a general advantage against no adaptations. Here, the SCAM strategy performs slightly better (highest accuracy and precision) than the other adaptive methods for the lower number of parameter models (BallStick_r1, Tensor) while the AMWG method performs slightly better in the higher number of parameter models (NODDI, CHARMED_r1). Generally the performance differences in accuracy and precision between the adaptive methods are not large, and both the SCAM and AWG perform well. Given the high alround efficiency, accuracy and precision and the maintained ergodicity of the chain in the AMWG method, we selected this method to generate chains in the rest of this work.

**Figure 3:**
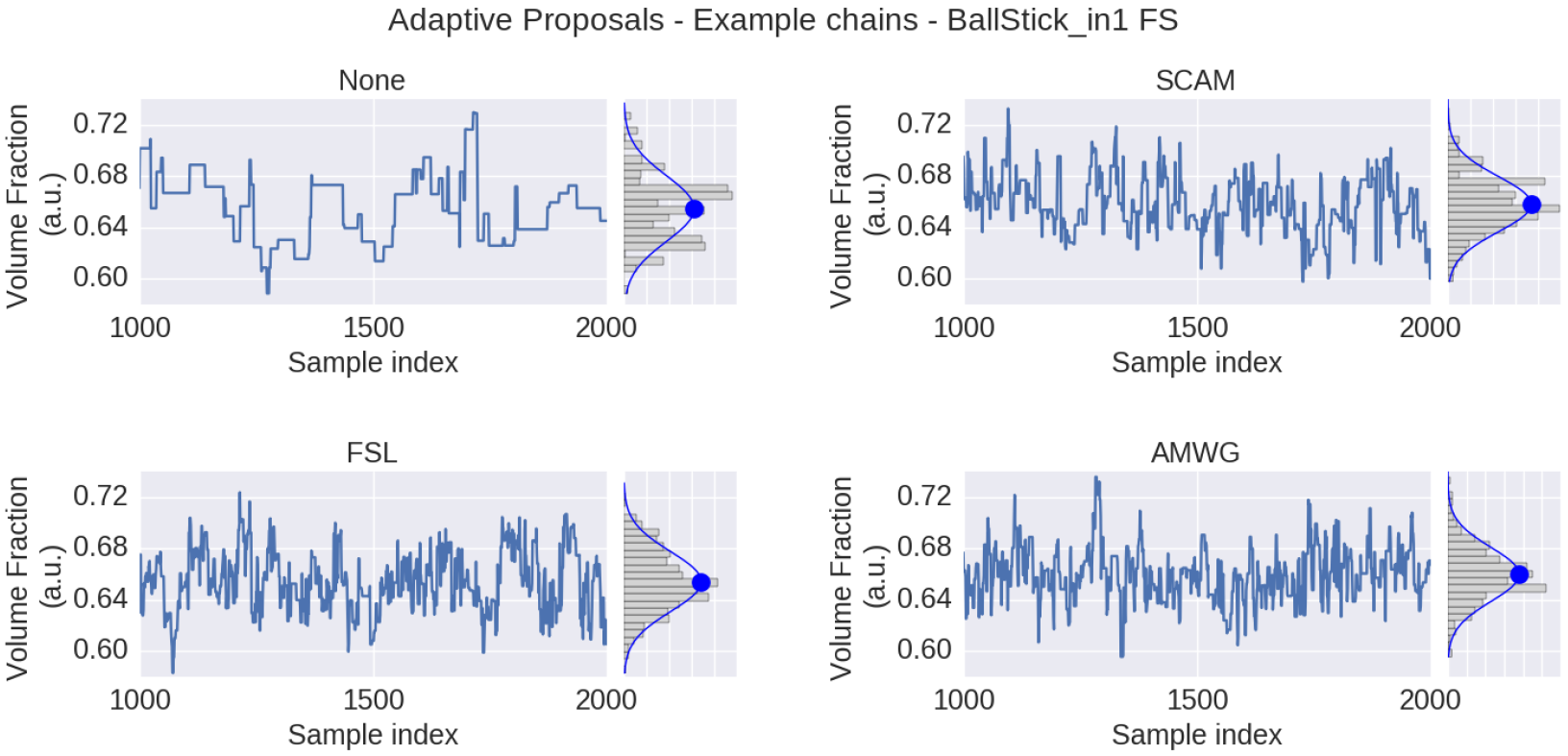
MCMC sample traces for the voxel indicated in figure 1, using Ball&Stick_in1 Fraction of Stick (FS), for no adaptive metropolis (None), the Single Component Adaptive Metropolis (SCAM), the FSL acceptance rate scaling (FSL) and Adaptive Metropolis-Within-Gibbs (AMWG) adaptive proposal methods. Results were computed with an initial proposal standard deviation of 0.25. A Gaussian distribution function was fitted to the samples, superimposed in blue on the sample histograms, with its mean indicated by the blue dot.

### 3.2 Burn-in

Figure 6 shows a comparison of mean and standard deviation estimates over 10,000 samples (no thinning), between sampling started from the default starting point (table 4) and from a Maximum Likelihood Estimator starting point, over an increasing length of burn-in. When started from a default starting point, the chains of most voxels will have converged to their stationary distribution after a burn-in of about 3000 samples. When started from an MLE starting point, the chain starts from a point in the stationary distribution and no burn-in is necessary. Starting from an MLE starting point has the additional advantage of removing salt-and pepper-like noise from the mean and std. maps. For example, even after a burn-in of 3000 samples, there are still some voxels in the default starting point maps that have not converged yet. Burn-in also seems to have a greater impact on the standard deviation estimates than it does on the mean estimates. After a burn-in of 1000 samples, the means of the default starting point maps seem to have converged, while the many of the standard deviations clearly have not. in contrast, stable standard deviation estimates are obtained from the MLE initialized chain even without burn-in.

**Figure 4:**
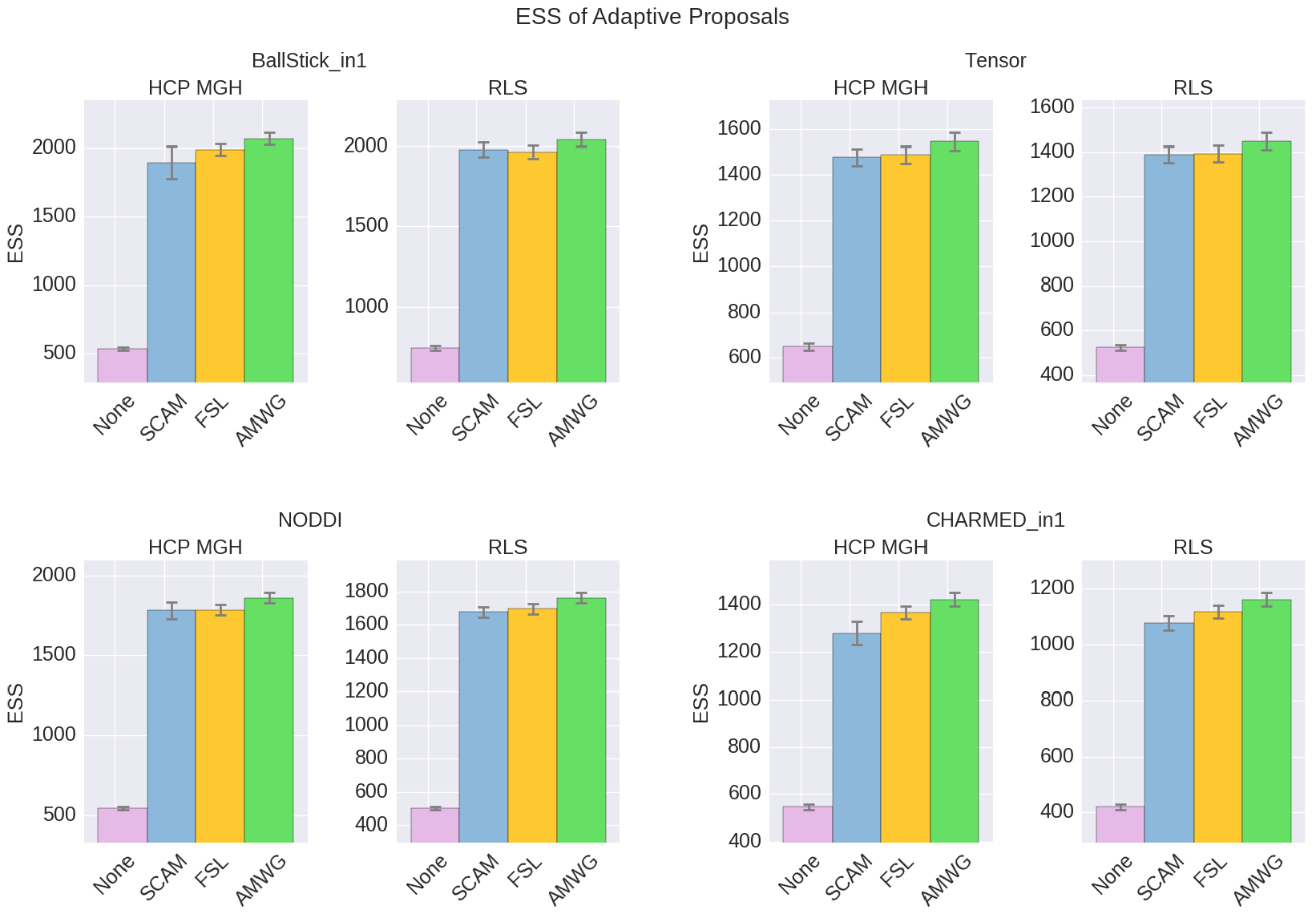
Estimated multivariate Effective Sample Size (ESS), for no adaptive metropolis (None), the Single Component Adaptive Metropolis (SCAM), the FSL acceptance rate scaling (FSL) and Adaptive Metropolis-Within-Gibbs (AMWG) adaptive proposal methods. Whiskers show the standard error of the mean computed over 10 repeats. Results are over 10000 samples, with a burn-in of 1000 samples, without thinning.

To illustrate this on a single chain basis, in figure 7 we plot the first 1000 samples of an MCMC run of the Ball&Stick_in1 and NODDI model using the MLE starting point and two random volume fractions as a starting point, on the left, and (on the right) the effect of discarding the first *z* samples when computing the posterior mean and standard deviation (with statistics over 1000 samples, after the burn-in *z*). The sampling trace shows how the sampler moves through the parameter space before converging to the stationary distribution. Interestingly, all points first seem to move toward an intra-axonal volume fraction of zero, before moving up again. This is probably caused by a misalignment of the model orientation with the data’s diffusion orientation, making the intra-axonal volume less likely. Only after a correct orientation of the model, the volume fraction can go up again. The plots on the right of the figure show the convergence of the mean and standard deviation with an increased burn-in length. These plots again show that, when started from the MLE, no burn-in is needed, while starting from another position some burn-in is required for the chains to converge.

**Figure 5:**
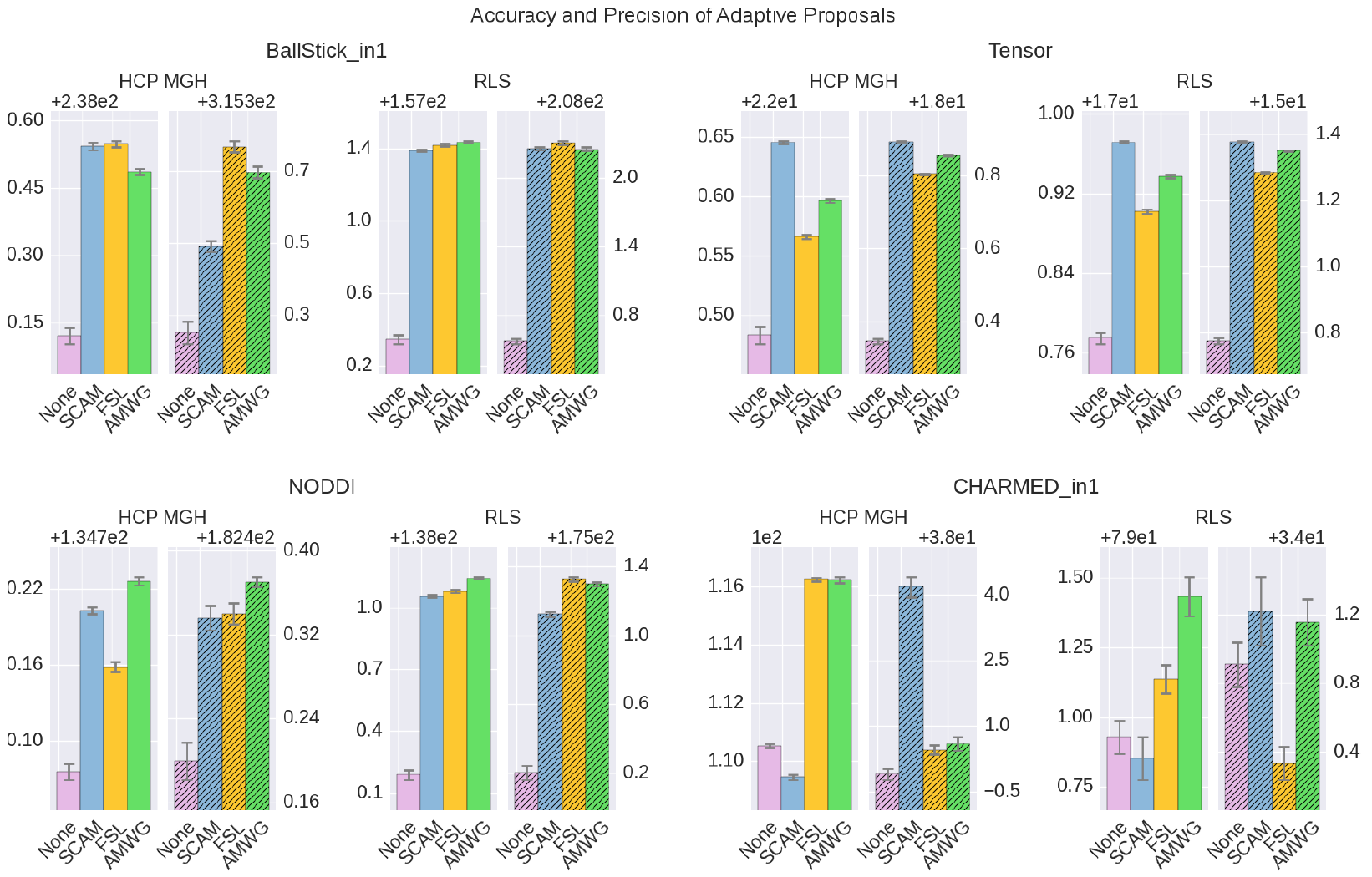
Estimated accuracy (left plots) and precision (right, shaded, plots), for no adaptive metropolis (None), the Single Component Adaptive Metropolis (SCAM), the FSL acceptance rate scaling (FSL) and Adaptive Metropolis-Within-Gibbs (AMWG) adaptive proposal methods. The results are averaged over 10000 voxels and 10 trials, the whiskers show the standard error of the mean computed over the 10 trials. Results are over 10000 samples, with a burn-in of 1000 samples, without thinning.

**Figure 6:**
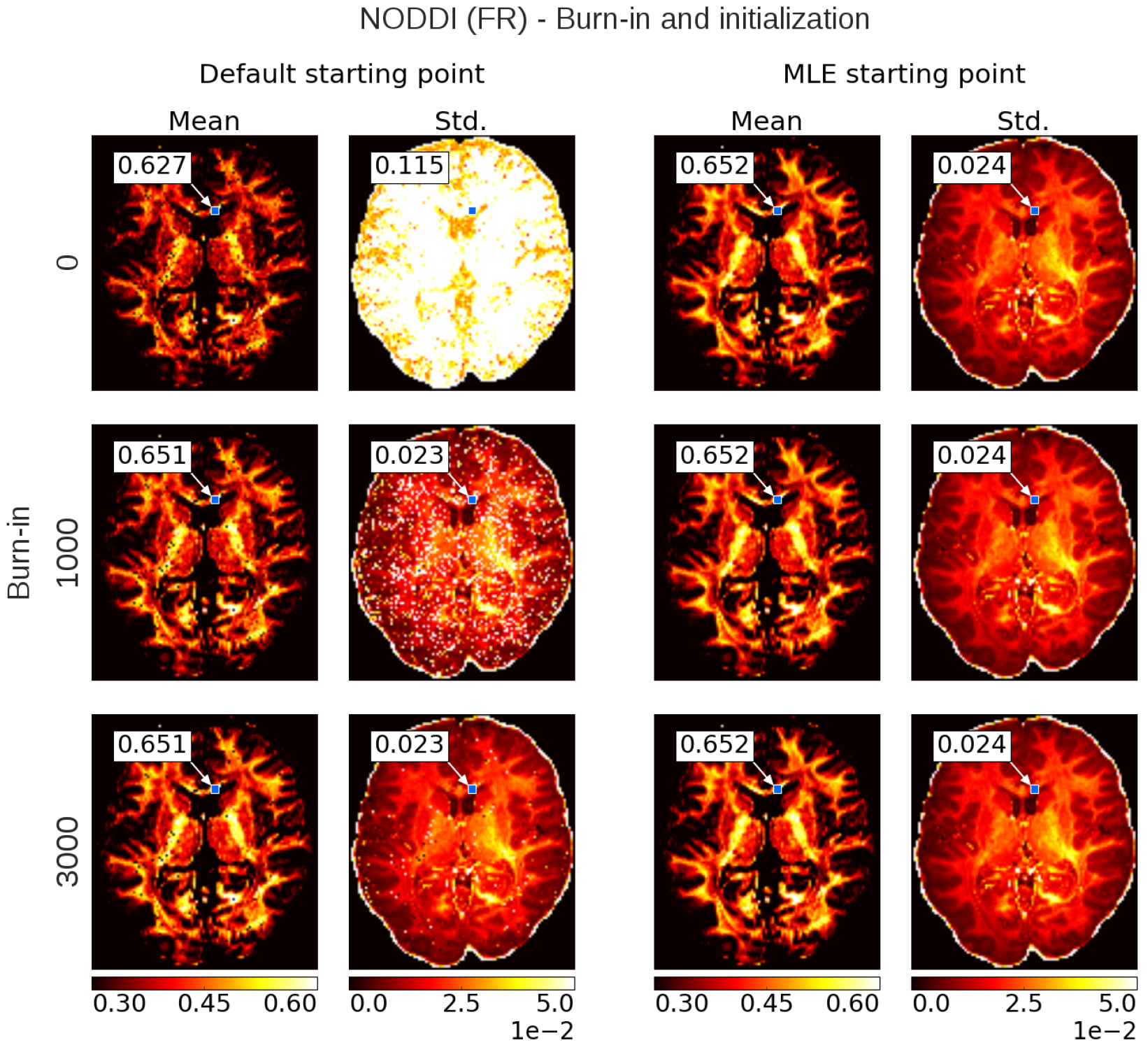
Burnin demonstration and chain initialization using NODDI Fraction of Restricted (FR). On the left, the posterior mean and standard deviation (std.) maps when sampling NODDI from the MDT default starting point, on the right the mean and std. maps when sampling NODDI using a Maximum Likelihood Estimator (MLE) as starting point. The rows show the effect of discarding the first *z* ∈ {0,1000, 3000} samples as burn-in before the mean and std. estimation. Statistics are without thinning and over 10,000 samples after *z*. The value insets show the mean and standard deviation value from a Gaussian fit to the sampling chain for the indicated voxel.

### 3.3 Thinning

Thinning of sampler chains has theoretically been shown to reduce the accuracy of posterior analyses (Geyer, 1991; MacEachern & Berliner, 1994; Link & Eaton, 2012), and empirical evidence has been provided for the limited usefulness of thinning Link2012 Here we will show some empirical results of thinning applied to diffusion MRI modeling. Figure 8 shows the effect of thinning on the variability of the returned sampling trace and on the estimates of the mean and standard deviation. The sampling trace shows that the chains produce roughly the same distribution, while with increased thinning many more samples are required (*k* times more samples, for a thinning of *k*). Comparing the effect of thinning on the mean and standard deviation shows that, as predicted by theory, there is less variance in the estimates when using more samples as compared to thinning the samples. Results also show that 1000 samples without thinning may not be enough for a stable estimates and more samples are required. Yet in accordance with theory, instead of thinning the chain, results indicate that just using more samples (e.g. all 1000 · *k* samples instead a thinning of *k*) is preferred.

**Figure 7:**
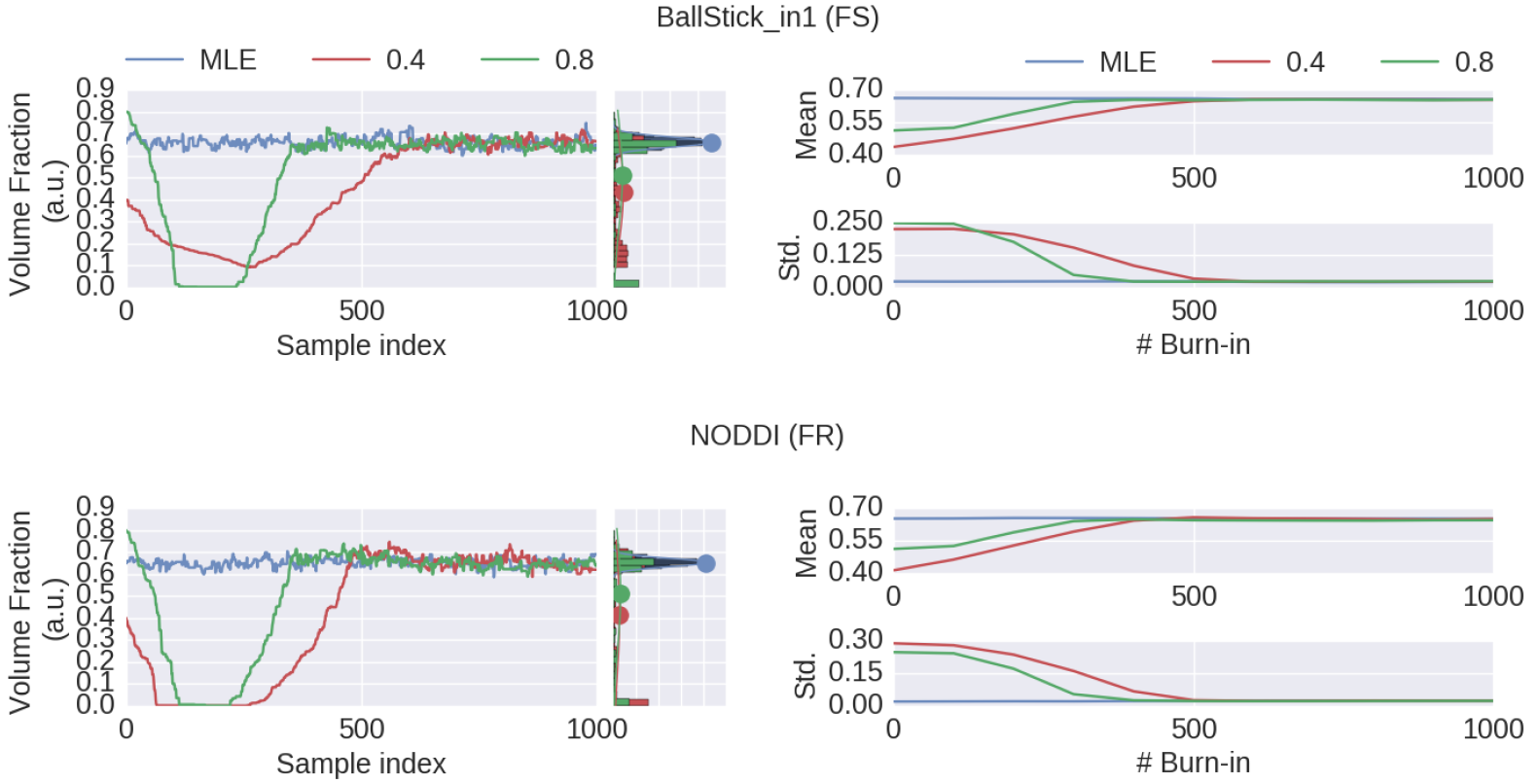
MCMC chains and burn-in results of a single voxel (the voxel indicated in 6). On the left, the sampling trace when starting at the MLE or two default points, with (only) a varying volume fraction. Chain histograms and Gaussian fits as before. On the right, mean and standard deviation computed over 1000 samples with increasing burn-in.

### 3.4 Minimum number of samples

Using multivariate ESS theory we determined, a priori, per model, the number of actual samples needed to generate a sufficient number of effective samples (the effective sample size or ESS) to approximate the underlying posterior density within a 95% confidence region and with a 90% relative precision. Figure 9 shows an estimate on the number of actual samples needed to reach this desired ESS, on average over a white matter mask. In general, the sampling requirements do not depend on the acquisition table, with similar numbers of samples needed for the HCP MGH and RLS protocols. The exception to the rule is in the CHARMED_r1 model where we need about double the number of samples for the RLS acquisition protocol compared to the HCP MGH protocol. This is probably due to the limited suitability of the RLS dataset for the CHARMED model as it requires high maximum b-values which the RLS protocol does not contain. Table 6 summarizes the estimated minimum sample requirements, together with the required ESS and the number of estimated parameters in each model. In general, models with more parameters need more actual samples to reach the same confidence and precision, although the Tensor model with seven parameters requires less samples than the NODDI model with six parameters. This is probably related to the higher complexity (nonlinear parameter inter-dependencies) of the NODDI model compared to the Tensor model. Interestingly, the required ESS to reach the desired confidence and precision is very similar, at about 2200, for all models (although the numbers of actual samples needed to realize this are different). As an estimate of computation times, table 7 shows runtime statistics for sampling the recommended number of samples for HCP MGH dataset and a RLS dataset using a AMD Fury X graphics card. This shows that although 4 to 7 hours are needed to sample more complex CHARMED and NODDI models on the very large HCP MGH dataset (with 552 volumes), gpu accelerated implementation can provide full posterior sampling of diffusion microstructure models over whole brain datasets in reasonable time on a standard graphics card. On the more clinicaly feasible RLS protocol (134 volumes) whole brain sampling of Tensor and Ball&Stick models can be performed within 20 minutes and NODDI within an hour.

**Figure 8:**
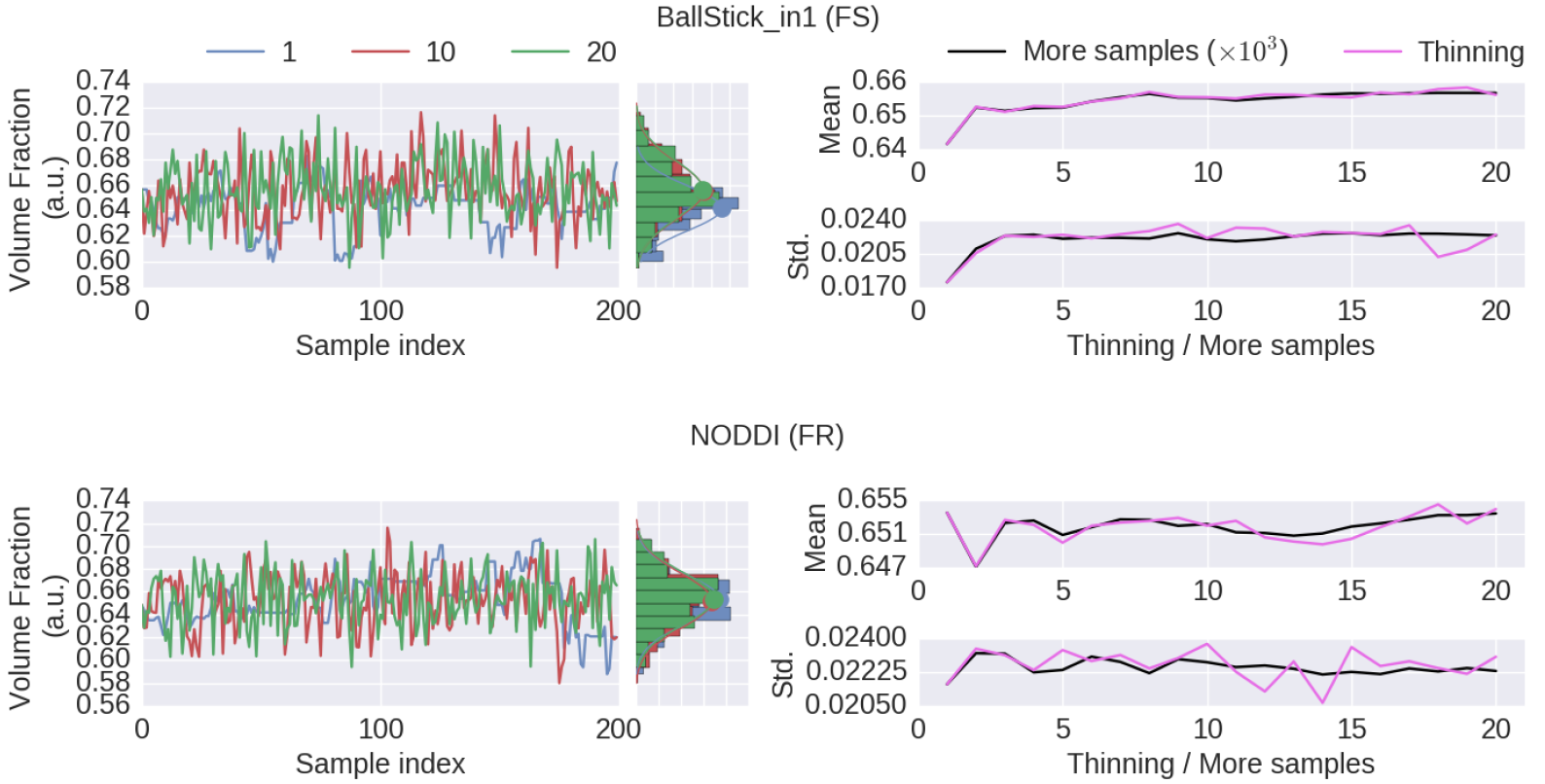
Thinning results of a single voxel (the voxel indicated in 6). On the left, sample traces for the returned samples after a thinning of 1 (no thinning), a thinning of 10 and of 20, with their corresponding histograms. On the right, a comparison of the posterior mean and standard deviation when thinning the chain or when using more samples. When thinning, 1000 · *k* samples are generated of which only every *k*th sample is used (so always 1000 samples are used). When using more samples, all 1000 · *k* samples are used, without thinning. Results are without burn-in and started from a maximum likelihood estimator.

**Figure 9:**
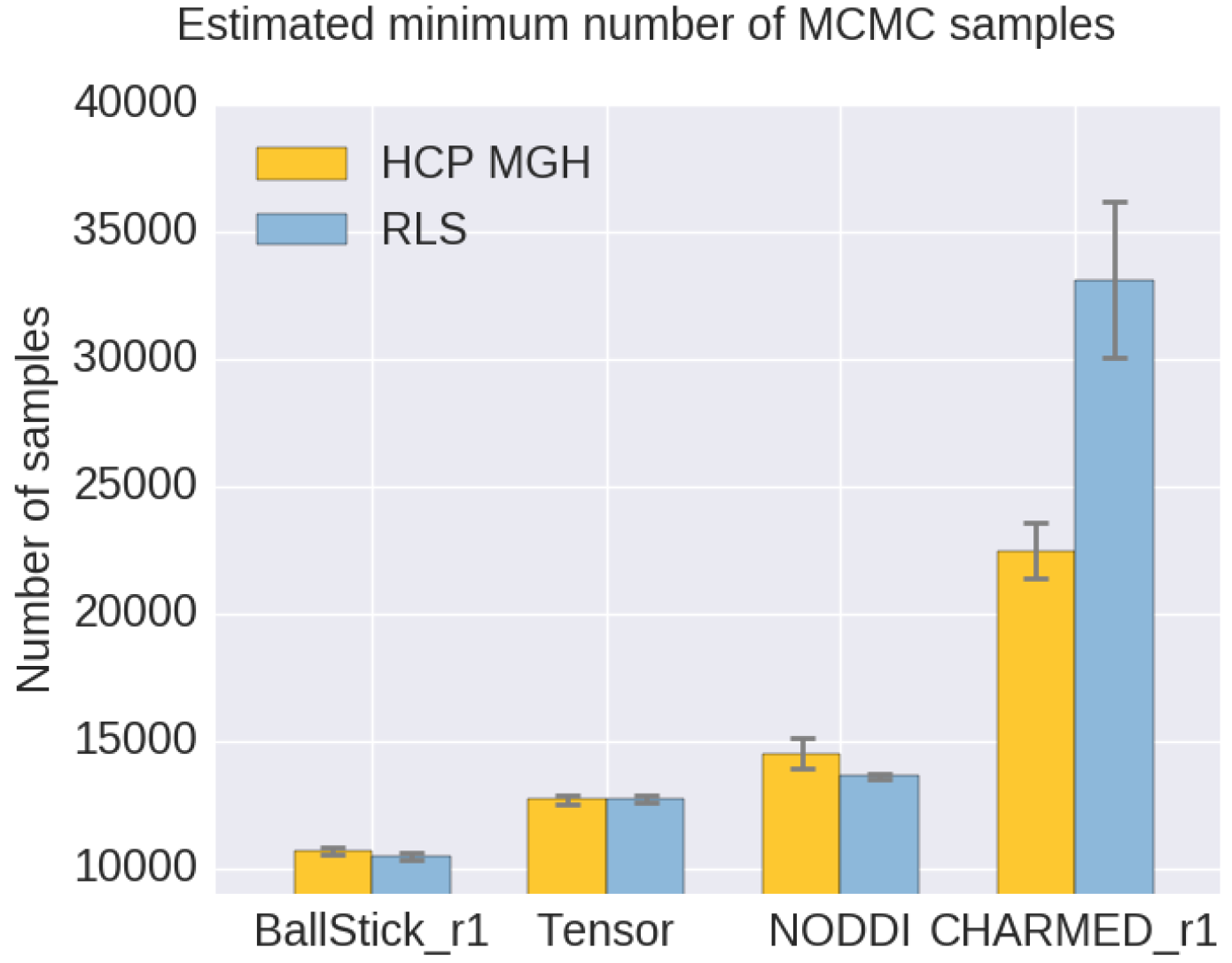
Estimates on the number of samples needed per model, to reach, when averaged over the white matter, a 95% confidence region with a 90% relative precision. Results are shown for both an HCP MGH and RLS acquisition table. Whiskers show the standard error of the mean over 10 subjects.

## 4 Discussion

Using an efficient GPU based implementation, we show that run times can be removed as a prohibitive constraint for sampling of diffusion multicompartment models, achieving whole brain sampling in under an hour for typical datasets and most common dMRI models. Newer generations of graphics cards are likely to reduce these times even further. Using this implementation, we investigated the use of adaptive MCMC algorithms, burn-in, initialization and thinning. We finally applied the theory of Effective Sample Size to diffusion multi-compartment models as a way of determining a sufficient number of samples for a given model and dataset.

**Table 6:** Estimates on the number of samples needed per model, to reach, when averaged over the white matter, a 95% confidence region with a 90% relative precision. While the required ESS can be determined a priori, the inherent model complexity determines how many samples are needed to reach that ESS.

| **Model** | **Number of parameters** | **Required ESS** | **Required nr. of samples** |
| --- | --- | --- | --- |
| BallStick_r1 | 4 | 2108 | 11000 |
| NODDI | 6 | 2177 | 15000 |
| Tensor | 7 | 2192 | 13000 |
| CHARMED_r1 | 11 | 2208 | 24000 |

**Table 7:** Runtime statistics (hh:mm:ss) for MCMC sampling the estimated minimum number of samples (no burn-in, no-thinning) of various models, using a single HCP MGH (552 volumes) and single RLS (134 volumes) dataset. Runtimes include prior model optimization using the Powell routine. Statistics are over a BET generated whole brain mask which include 410,000 voxels for HCP MGH and 204,993 voxels for RLS. Results are computed using a single AMD Fury X graphics card.

| **Model** | **Number of samples** | **HCP MGH** | **RLS** |
| --- | --- | --- | --- |
| Ball&Stick_in1 | 11000 | 00:47:18 | 00:12:56 |
| Tensor | 13000 | 00:48:53 | 00:19:55 |
| NODDI | 15000 | 04:20:03 | 00:54:43 |
| CHARMED_in1 | 24000 | 07:15:13 | 01:54:51 |

### 4.1 Adaptive MCMC

The use of adaptive MCMC algorithms increases both the estimated multivariate effective sample sizes as well as the accuracy and precision of posterior mean estimates. The Adaptive Metropolis-Within-Gibbs (*AMWG*) outperforms other proposal adaptation methods in terms of multivariate Effective Sample Size (ESS). In accuracy and precision the AMWG method performs higher for the NODDI and CHARMED in1 models, but (slightly) lower for the Ball&Stick_in1 and Tensor models where, respectively, the FSL and the Single Component Adaptive Metropolis (*SCAM*) methods perform better. Performance of the fixed proposal method could, in theory, be increased to the same levels as the adaptive methods by manual calibration, which could also slightly decrease the chain’s autocorrelation compared to adaptive proposals. Since this is model and data (and thus voxel) dependent, manual tuning could be very burdensome and unpractical. This work covers only variations of the single component Random Walk Metropolis, which have the advantage of high efficiency sampling with relatively general model-unspecific proposals. Future work could focus on MCMC algorithms which allow for block-updates of correlated parameters, or could investigate different proposal schemes altogether such as Component-wise Hit-And-Run Metropolis (Turchin, 1971; Smith, 1984), Multiple-Try Metropolis (Liu et al., 2000) and/or No-U-Turn sampler (Hoffman & Gelman, 2011).

### 4.2 Burn-in

When starting from an arbitrary position, burn-in is advisable to reduce possible bias due to (possibly) low probability starting positions. Burn-in should ideally be considered post-sampling, since it is difficult to know a priori the time needed for the chain to converge and, due to randomness, past convergence rates provide no guarantee for the future. This is why common practice dictates a relatively large number of burn-in samples which guarantees convergence in most cases.

While not harmful, burn-in is generally unnecessary and inefficient if the starting point is part of the stationary distribution of the Markov chain, which can, for example, be achieved by taking a Maximum Likelihood Estimator (MLE) as starting point. Even when starting from an MLE, a small burn-in of about 100 to 200 samples could be considered to remove correlations with the starting position. Additionally, when using adaptive proposal methods, a small burn-in could be considered to let the adaptation algorithm adapt the proposal distribution before sampling, slightly increasing the effective sample size of the chain.

### 4.3 Thinning

Already on theoretical grounds, thinning is not recommended and considered as often unnecessary, always inefficient and reducing the precision of posterior estimates (Link & Eaton, 2012; MacEachern & Berliner, 1994; Jackman, 2009; Geyer, 1991; Christensen et al., 2010). Illustrations based on the the Ball&Stick_in1 and NODDI model show that, with or without thinning, the posterior distribution is approximated about equally, while thinning needs *k* times more samples (for a thinning of *k*). Results did show a convergence of mean and standard deviation estimates with an increased thinning, but these results are easily duplicated by incorporating not only the thinned samples but also the non-thinned samples in the statistical estimates (the ‘more samples’ strategy). Furthermore, using more samples instead of thinning provides estimates with a higher precision, as illustrated by the higher variability of the thinned estimates compared to the estimates with more samples (Figure 8, right). One legitimate reason for thinning is that, with independent samples, one can approximate the precision of an MCMC approximation (Link & Eaton, 2012). That is, it allows for more accurate assessment of the standard error of an MCMC estimate like the posterior mean. However, even in that case, thinning must be applied post-hoc, otherwise the precision of the mean itself will be reduced if computed from only the thinned samples. Furthermore, we are often more interested in the variability of the posterior distribution (which can be provided by e.g. the standard deviation) than in the precision of the posterior mean estimate. Another legitimate reason for considering thinning is hardware limitations, such as sampling post-processing time and storage space. However, barring such limitations, avoiding thinning of chains is far more efficient in providing high precision in posterior estimates.

### 4.4 Number of samples

The issue of the number of samples needed in a chain is often somewhat enigmatic and arbitrary. A common perception is that the number should be ‘high’, rather too high than too low. Multivariate Effective Sample Size (ESS) theory provides a theoretical lower bound on the number of effective samples needed to approximate the posterior, based on a desired confidence level and precision. How many MCMC samples are required to reach that target effective sample size is then dependent on the data, the model and the MCMC algorithm. We show that the dependency on the data seems to be low for diffusion microstructure models, considering the approximately equal sampling requirements using two different datasets for all but the CHARMED_in1 model. For that model, the sampling requirements are almost twice as high for the RLS as for the MGH dataset, probably because the RLS acquisition is not suitable for the CHARMED_-in1 model. The dependency on the model is higher, showing that more complex models seem to need more actual samples to reach the target ESS. Interestingly, we show that for a 95% confidence region (*α* = 0.05) with a 90% precision (*ϵ* = 0.1) the target ESS is about 2200 for all models. This sets an informed relatively general target for the amount of samples required in sampling diffusion microstuctural models, which scales the number of actual samples with the complexity of the model, data and the performance of the MCMC algorithm. This also means that MCMC algorithms that can generate effective samples more efficiently (such as the AMWG) can reduce the number of samples needed to reach the same confidence levels, reducing run-time.

## 5 Conclusions and recommendations

Considering the theoretical soundness and its general robust performance, we advise the Adaptive Metropolis-Within-Gibbs (AMWG) algorithm for efficient and robust sampling of diffusion MRI models. We further recommend initializing the sampler with a maximum likelihood estimator obtained from, for example, non-linear optimization, in which case 100 to 200 samples are sufficient as a burn-in. Thinning is unnecessary unless there are memory or hard disk constraints or a strong reliance on posterior estimates that require uncorrelated samples. As a relatively general target for the number of samples, we recommend 2200 multivariate effective samples. The amount of actual MCMC samples required to achieve this is algorithm and model dependent and can be investigated in a pre-study, with numbers for common dMRI models reported here as an indication.

## 6 Acknowledgements

RLH and AR were supported by an ERC Starting Grant (MULTICONNECT, #639938), AR was additionally supported by a Dutch science foundation (NWO) VIDI Grant (#14637). Data collection and sharing for this project was provided, in part, by the MGH-USC Human Connectome Project (HCP; Principal Investigators: Bruce Rosen, M.D., Ph.D., Arthur W. Toga, Ph.D., Van J. Weeden, MD). HCP funding was provided by the National Institute of Dental and Craniofacial Research (NIDCR), the National Institute of Mental Health (NIMH), and the National Institute of Neurological Disorders and Stroke (NINDS). HCP data are disseminated by the Laboratory of Neuro Imaging at the University of California, Los Angeles. Collectively, the HCP is the result of efforts of co-investigators from the University of California, Los Angeles, Martinos Center for Biomedical Imaging at Massachusetts General Hospital (MGH), Washington University, and the University of Minnesota. Data collection and sharing for this project was provided, in part, by the Rhineland Study www.rheinland-studie.de, Principal Investigator: Monique M.B. Breteler, M.D., Ph.D.; German Center for Neurodegenerative Diseases (DZNE), Bonn).

## References

Alexander, D. C. (2008). A general framework for experiment design in diffusion MRI and its application in measuring direct tissue-microstructure features. Magnetic resonance in medicine, 60, 439–48. URL: http://www.ncbi.nlm.nih.gov/pubmed/18666109. doi: 10.1002/mrm.21646.

Alexander, D. C. (2009). Modelling, Fitting and Sampling in Diffusion MRI. In L. D., & W. J. (Eds.), Visualization and Processing of Tensor Fields. Mathematics and Visualization. (pp. 3–20). Springer, Berlin, Heidelberg. URL: http://link.springer.com/10.1007/978-3-540-88378-4{_}1. doi: 10.1007/978-3-540-88378-4_1.

Alexander, D. C., Hubbard, P. L., Hall, M. G., Moore, E. A., Ptito, M., Parker, G. J. M., & Dyrby, T. B. (2010). Orientationally invariant indices of axon diameter and density from diffusion MRI. NeuroImage, 52, 1374–1389. URL: http://dx.doi.org/10.1016Zj.neuroimage.2010.05.043. doi:10.1016/j.neuroimage.2010.05.043.

Assaf, Y., Alexander, D. C., Jones, D. K., Bizzi, A., Behrens, T. E. J., Clark, C. A., Cohen, Y., Dyrby, T. B., Huppi, P. S., Knoesche, T. R., LeBihan, D., Parker, G. J. M., Poupon, C., Anaby, D., Anwander, A., Bar, L., Barazany, D., Blumenfeld-Katzir, T., De-Santis, S., Duclap, D., Figini, M., Fischi, E., Guevara, P., Hubbard, P., Hofstetter, S., Jbabdi, S., Kunz, N., Lazeyras, F., Lebois, A., Liptrot, M. G., Lundell, H., Man-gin cedil;ois, J. F., Dominguez, D. M., Morozov, D., Schreiber, J., Se-unarine, K., Nava, S., Riffert, T., Sasson, E., Schmitt, B., Shemesh, N., Sotiropoulos, S. N., Tavor, I., Zhang, H., & Zhou, F. L. (2013). The CONNECT project: Combining macro- and micro-structure. NeuroImage, 80, 273–282. URL: http://dx.doi.org/10.1016/j.neuroimage.2013.05.055. doi:10.1016/j.neuroimage.2013.05.055.

Assaf, Y., & Basser, P. J. (2005). Composite hindered and restricted model of diffusion (CHARMED) MR imaging of the human brain. NeuroImage, 27, 48–58. doi:10.1016/j.neuroimage.2005.03.042.

Assaf, Y., Blumenfeld-Katzir, T., Yovel, Y., & Basser, P. J. (2008). AxCaliber: A method for measuring axon diameter distribution from diffusion MRI. Magnetic Resonance in Medicine, 59, 1347–1354. URL: http://www.ncbi.nlm.nih.gov/pubmed/185067999. doi:10.1002/mrm.21577. arXiv:15334406.

Assaf, Y., Freidlin, R. Z., Rohde, G. K., & Basser, P. J. (2004). New modeling and experimental framework to characterize hindered and restricted water diffusion in brain white matter. Magnetic Resonance in Medicine, 52, 965–978. URL: http://www.ncbi.nlm.nih.gov/pubmed/15508168. doi:10.10 02/mrm.20274.

Basser, P. J., Mattiello, J., & LeBihan, D. (1994). MR diffusion tensor spectroscopy and imaging. Biophysical journal, 66, 259–67. URL: http://www.pubmedcentral.nih.gov/articlerender.fcgi?artid=1275686{&}tool=pmcentrez{&}rendertype=abstract. doi:10.1016/S0006-3495(94)80775-1.

Behrens, T. E. J., Woolrich, M. W., Jenkinson, M., Johansen-Berg, H., Nunes, R. G., Clare, S., Matthews, P. M., Brady, J. M., & Smith, S. M. (2003). Characterization and Propagation of Uncertainty in Diffusion-Weighted MR Imaging. Magnetic Resonance in Medicine, 50, 1077–1088. URL: http://www.ncbi.nlm.nih.gov/pubmed/14587019. doi:10.1002/mrm.10609.

Chib, S., & Greenberg, E. (1995). Understanding the Metropolis-Hastings Algorithm. The American Statistician, 49, 327–335.

Christensen, R., Johnson, W., Branscum, A., & Hanson, T. E. (2010). Bayesian Ideas and Data Analysis: An Introduction for Scientists and Statisticians.

Daducci, A., Canales-Rodríguez, E. J., Zhang, H., Dyrby, T. B., Alexander, D. C., & Thiran, J. P. (2015). Accelerated Microstructure Imaging via Convex Optimization (AMICO) from diffusion MRI data. NeuroImage, 105, 32–44. URL: http://www.ncbi.nlm.nih.gov/pubmed/25462697 http://dx.doi.org/10.1016/j.neuroimage.2014.10.026. doi:10.1016/j.neuroimage.2014.10.026.

De Santis, S., Assaf, Y., Evans, C. J., & Jones, D. K. (2014a). Improved precision in CHARMED assessment of white matter through sampling scheme optimization and model parsimony testing. Magnetic Resonance in Medicine, 71, 661–671. URL: http://www.ncbi.nlm.nih.gov/pubmed/23475834. doi:10.1002/mrm.24717.

De Santis, S., Drakesmith, M., Bells, S., Assaf, Y., & Jones, D. K. (2014b). Why diffusion tensor MRI does well only some of the time: Variance and covariance of white matter tissue microstructure attributes in the living human brain. NeuroImage, 89, 35–44. URL: http://dx.doi.org/10.1016Zj.neuroimage.2013.12.003 http://www.ncbi.nlm.nih.gov/pubmed/24342225. doi:10.1016/j.neuroimage.2013.12.003.

De Santis, S., Jones, D. K., & Roebroeck, A. (2016). Including diffusion time dependence in the extra-axonal space improves in vivo estimates of axonal diameter and density in human white matter. NeuroImage, 130, 91–103. URL: http://dx.doi.org/10.1016/j.neuroimage.2016.01.047. doi:10.1016/j.neuroimage.2016.01.047.

Dietrich, O., Raya, J. G., Reeder, S. B., Reiser, M. F., & Schoenberg, S. O. (2007). Measurement of signal-to-noise ratios in MR images: Influence of multichannel coils, parallel imaging, and reconstruction filters. Journal of Magnetic Resonance Imaging, 26,375–385. doi:10.1002/jmri.20969.

Fieremans, E., Benitez, A., Jensen, J. H., Falangola, M. F., Tabesh, A., Dear-dorff, R. L., Spampinato, M. V. S., Babb, J. S., Novikov, D. S., Ferris, S. H., & Helpern, J. A. (2013). Novel white matter tract integrity metrics sensitive to Alzheimer disease progression. American Journal of Neuroradiology, 34, 2105–2112. doi:10.317 4/ajnr.A3553.

Gelman, A., Carlin, J. B., Stern, H. S., Dunson, D. B., Vehtari, A., & Rubin, D. B. (2013). Bayesian Data Analysis. CRC Press.

Gelman, A., Roberts, G., & Gilks, W. (1996). Efficient Metropolis Jumping Rules. Bayesian statistics 5, (pp. 599–608).

Geman, S., & Geman, D. (1984). Stochastic Relaxation, Gibbs Distributions, and the Bayesian Restoration of Images. IEEE Transactions on Pattern Analysis and Machine Intelligence, PAMI-6, 721–741. doi:10.1109/TPAMI.1984.4767596.

Geyer, C. J. (1991). Markov Chain Monte Carlo Maximum Likelihood. Computing Science and Statistics: Proceedings of the 23rd Symposium on the Interface, (pp. 156–163).

Gong, L., & Flegal, J. M. (2016). A Practical Sequential Stopping Rule for High-Dimensional Markov Chain Monte Carlo. Journal of Computational and Graphical Statistics, 25, 684–700. URL: http://www.tandfonline.com/doi/full/10.1080/10618600.2015.1044092. doi:10.1080/10618600.2015.1044092. arXiv:1403.5536.

Haario, H., Saksman, E., & Tamminen, J. (2005). Componentwise adaptation for high dimensional MCMC. Computational Statistics, 20, 265–273. doi:10.1007/BF02789703.

Harms, R. L., Fritz, F. J., Tobisch, A., Goebel, R., & Roebroeck, A. (2017). Robust and fast nonlinear optimization of diffusion MRI microstructure models. NeuroImage, 155, 82–96. URL: http://dx.doi.org/10.1016/j.neuroimage.2017.04.064 http://linkinghub.elsevier.com/retrieve/pii/S1053811917303178. doi:10.1016/j.neuroimage.2017.04.064.

Hastings, W. K. (1970). Monte Carlo Sampling Methods Using Markov Chains and Their Applications. Biometrika, 57, 97. URL: http://www.jstor.org/stable/2334940?origin=crossref. doi:10.2307/2334940.

Hoffman, M. D., & Gelman, A. (2011). The No-U-Turn Sampler: Adaptively Setting Path Lengths in Hamiltonian Monte Carlo. Journal of Machine Learning Research, 15, 1593–1623. URL: http://arxiv.org/abs/1111.4246.arXiv:1111.4246.

Jackman, S. (2009). Bayesian Analysis for the Social Sciences. Chichester, UK: John Wiley & Sons, Ltd. URL: http://doi.wiley.com/10.1002/9780470686621. doi:10.1002/9780470686621.

Jelescu, I. O., Veraart, J., Adisetiyo, V., Milla, S. S., Novikov, D. S., & Fieremans, E. (2015a). One diffusion acquisition and different white matter models: How does microstructure change in human early development based on WMTI and NODDI? NeuroImage, 107, 242–256. URL: http://dx.doi.org/10.1016/j.neuroimage.2014.12.009 http://www.ncbi.nlm.nih.gov/pubmed/25498427. doi:10.1016/j.neuroimage.2014.12.009.

Jelescu, I. O., Veraart, J., Fieremans, E., & Novikov, D. S. (2015b). Caveats of non-linear fitting to brain tissue models of diffusion. In ISMRM 2015 (p. 88040). volume 23.

Jelescu, I. O., Veraart, J., Fieremans, E., & Novikov, D. S. (2016). Degeneracy in model parameter estimation for multi-compartmental diffusion in neuronal tissue. NMR in Biomedicine, 29, 33–47. URL: http://www.ncbi.nlm.nih.gov/pubmed/26615981. doi:10.1002/nbm.3450.

Johnson, A. A., Jones, G. L., & Neath, R. C. (2013). Component-Wise Markov Chain Monte Carlo: Uniform and Geometric Ergodicity under Mixing and Composition. Statistical Science, 28, 360–375. URL: http://projecteuclid.org/euclid.ss/1377696941. doi:10.1214/13-STS423. arXiv:arXiv:0903.0664v6.

Kass, R. E., Carlin, B. P., Gelman, A., & Neal, R. M. (1998). Markov Chain Monte Carlo in Practice: A Roundtable Discussion. The American Statistician, 52, 93. URL: http://www.jstor.org/stable/2685466?origin=crossref. doi:10.2307/2685466.

Le Bihan, D., Breton, E., Lallemand, D., Grenier, P., Cabanis, E., & Laval-Jeantet, M. (1986). MR imaging of intravoxel incoherent motions: application to diffusion and perfusion in neurologic disorders. Radiology, 161, 401–407. URL: http://pubs.rsna.org/doi/10.1148/radiology.161.2.3763909. doi:10.1148/radiology.161.2.3763909.

Link, W. A., & Eaton, M. J. (2012). On thinning of chains in MCMC. Methods in Ecology and Evolution, 3, 112–115. doi:10.1111/j.2041-210X.2011.00131.x.

Liu, J. S. (2004). Monte Carlo Strategies in Scientific Computing. Springer Series in Statistics. New York, NY: Springer New York. URL: http://link.springer.com/10.1007/978-0-387-76371-2. doi:10.1007/978-0-387-76371-2.

Liu, J. S., Liang, F., & Wong, W. H. (2000). The Multiple-Try Method and Local Optimization in Metropolis Sampling. Journal of the American Statistical Association, 95, 121. URL: http://www.jstor.org/stable/2669532?origin=crossref. doi:10.2307/2669532.

MacEachern, S. N., & Berliner, L. M. (1994). Subsampling the Gibbs Sampler. The American Statistician, 48, 188. URL: http://www.jstor.org/stable/2684714?origin=crossref. doi:10.2307/2684714.

Martino, L., Elvira, V., & Louzada, F. (2017). Effective sample size for importance sampling based on discrepancy measures. Signal Processing, 131, 386–401. doi:10.1016/j.sigpro.2016.08.025. arXiv:1602.03572.

Metropolis, N., Rosenbluth, A. W., Rosenbluth, M. N., Teller, A. H., & Teller, E. (1953). Equation of State Calculations by Fast Computing Machines. The Journal of Chemical Physics, 21, 1087–1092. URL: http://aip.scitation.org/doi/10.1063/!.1699114. doi:10.1063/1.1699114. arXiv:5744249209.

Moeller, S., Yacoub, E., Olman, C. A., Auerbach, E., Strupp, J., Harel, N., & Ugurbil, K. (2010). Multiband multislice GE-EPI at 7 tesla, with 16fold acceleration using partial parallel imaging with application to high spatial and temporal whole-brain FMRI. Magnetic Resonance in Medicine, 63,1144–1153. doi:10.1002/mrm.22361.

Muller, P. (1994). Metropolis based posterior integration schemes. Numerical Recipes in Fortran (2nd Edition),. URL: http://citeseerx.ist.psu.edu/viewdoc/summary?doi=10.1.1.55.3539.

Panagiotaki, E., Schneider, T., Siow, B., Hall, M. G., Lyth-goe, M. F., & Alexander, D. C. (2012). Compartment models of the diffusion MR signal in brain white matter: A taxonomy and comparison. NeuroImage, 59, 2241–2254. URL: http://dx.doi.org/10.1016/j.neuroimage.2011.09.081 http://www.ncbi.nlm.nih.gov/pubmed/22001791. doi:10.1016/j.neuroimage.2011.09.081.

van Ravenzwaaij, D., Cassey, P., & Brown, S. D. (2016). A simple introduction to Markov Chain Monte-Carlo sampling. Psychonomic Bulletin and Review, (pp. 1–12). URL: http://dx.doi.org/10.3758/s13423-016-1015-8. doi:10.3758/s13423-016-1015-8.

Robert, C. P. (2015). The Metropolis-Hastings algorithm. Technical Report Wiley StatsRef: Statistics Reference Online. URL: http://arxiv.org/abs/1504.01896. doi:10.1002/9781118445112.stat07834. arXiv:1504.01896.

Robert, C. P., & Casella, G. (2004). Monte Carlo Statistical Methods. Springer Texts in Statistics. New York, NY: Springer New York. URL: http://link.springer.com/10.1007/978-1-4757-4145-2. doi:10.1007/978-1-4757-4145-2.

Roberts, G. O., & Rosenthal, J. S. (2007). Coupling and ergodicity of adaptive Markov chain Monte Carlo algorithms. Journal of Applied Probability, 44, 458–475. URL: http://projecteuclid.org/euclid.jap/1183667414. doi:10.1239/jap/1183667414.

Roberts, G. O., & Rosenthal, J. S. (2009). Examples of adaptive MCMC. Journal of Computational and Graphical Statistics, 18,349–367. URL: http://probability.ca/jeff/ftpdir/adaptex.pdf. doi:10.1198/jcgs.2009.06134.

Santis, S. D., Assaf, Y., Jeurissen, B., Jones, D. K., & Roebroeck, A. (2016). T1 relaxometry of crossing fibres in the human brain. NeuroImage, 141, 133–142. URL: http://linkinghub.elsevier.com/retrieve/pii/S1053811916303445. doi:10.1016/j.neuroimage.2016.07.037.

Sherlock, C., Fearnhead, P., & Roberts, G. O. (2010). The Random Walk Metropolis: Linking Theory and Practice Through a Case Study. Statistical Science, 25, 172–190. URL: http://projecteuclid.org/euclid.ss/1290175840. doi:10.1214/10-STS327. arXiv:arXiv:1011.6217v1.

Smith, R. L. (1984). Efficient Monte Carlo Procedures for Generating Points Uniformly Distributed over Bounded Regions. Operations Research, 32, 1296–1308. URL: http://pubsonline.informs.org/doi/abs/10.1287/opre.32.6.1296. doi:10.1287/opre.32.6.1296.

Smith, S. M. (2002). Fast robust automated brain extraction. Human Brain Mapping, 17,143–155. URL: http://www.ncbi.nlm.nih.gov/pubmed/12391568. doi:10.1002/hbm.10062.

Sotiropoulos, S. N., Behrens, T. E. J., & Jbabdi, S. (2012). Ball and rackets: Inferring fiber fanning from diffusion-weighted MRI. NeuroImage, 60, 1412–1425. URL: http://www.ncbi.nlm.nih.gov/pubmed/22270351. doi:10.1016/j.neuroimage.2012.01.056.

Sotiropoulos, S. N., Jbabdi, S., Andersson, J. L., Woolrich, M. W., Ugurbil, K., & Behrens, T. E. J. (2013). RubiX: Combining spatial resolutions for bayesian inference of crossing fibers in diffusion MRI. IEEE Transactions on Medical Imaging, 32, 969–982. URL: http://www.ncbi.nlm.nih.gov/pubmed/23362247. doi:10.1109/TMI.2012.2231873.

Tariq, M., Schneider, T., Alexander, D. C., Gandini Wheeler-Kingshott, C. A., & Zhang, H. (2016). Bingham-NODDI: Mapping anisotropic orientation dispersion of neurites using diffusion MRI. NeuroImage, 133, 207–223. URL: https://www.ncbi.nlm.nih.gov/pubmed/26826512. doi:10.1016/j.neuroimage.2016.01.046.

Turchin, V. F. (1971). On the Computation of Multidimensional Integrals by the Monte-Carlo Method. Theory of Probability & Its Applications, 16, 720–724. URL: http://epubs.siam.org/doi/10.1137/1116083. doi:10.1137/1116083.

Vats, D., Flegal, J. M., & Jones, G. L. (2015). Multivariate Output Analysis for Markov chain Monte Carlo. ArXiv e-prints, (pp. 1–52). URL: http://arxiv.org/abs/1512.07713.arXiv:1512.07713.

Xu, J., Moeller, S., Auerbach, E. J., Strupp, J., Smith, S. M., Feinberg, D. A., Yacoub, E., & Ugurbil, K. (2013). Evaluation of slice accelerations using multiband echo planar imaging at 3T. NeuroImage, 83,991–1001. doi:10.1016/j.neuroimage.2013.07.055.arXiv:NIHMS150003.

Zhang, H., Hubbard, P. L., Parker, G. J. M., & Alexander, D. C. (2011). Axon diameter mapping in the presence of orientation dispersion with diffusion MRI. NeuroImage, 56, 1301–1315. URL: http://dx.doi.org/10.1016/j.neuroimage.2011.01.084 http://www.ncbi.nlm.nih.gov/pubmed/21316474. doi:10.1016/j.neuroimage.2011.01.084.

Zhang, H., Schneider, T., Wheeler-Kingshott, C. A., & Alexander, D. C. (2012). NODDI: Practical in vivo neurite orientation dispersion and density imaging of the human brain. NeuroImage, 61, 1000–1016. URL: http://dx.doi.org/10.1016Zj.neuroimage.2012.03.072 http://www.ncbi.nlm.nih.gov/pubmed/22484410 http://ac.els-cdn.com/S1053811912003539/1-s2.0-S1053811912003539-main.pdf?{_}tid=55731f80-bac3-11e5-b1b1-00000aab0f26{&}acdnat=1452778562{_}8419fbce016e7afd001b1a2d30. doi:10.1016/j.neuroimage.2012.03.072.

